# Particulate matter composition drives differential molecular and morphological responses in lung epithelial cells

**DOI:** 10.1101/2023.05.17.541204

**Authors:** Sean M. Engels, Pratik Kamat, G. Stavros Pafilis, Yukang Li, Anshika Agrawal, Daniel J. Haller, Jude M. Phillip, Lydia M. Contreras

**Author notes:** **Corresponding Authors:** Lydia Contreras and Jude Phillip **Email:** and. **Author Contributions:** S.M.E., P.K., J.M.P., and L.M.C. designed the study. S.M.E., P.K., G.S.P., Y.L., A.A., and D.J.H. performed the experiments. S.M.E. and P.K. led analysis of the results. S.M.E. and P.K. wrote the manuscript. S.M.E., P.K., J.M.P., L.M.C., G.S.P., and D.J.H. edited the manuscript. L.M.C. and J.M.P. supervised the study. **Competing Interest Statement:** The authors declare that they have no competing interests.

## Abstract

Particulate matter (PM) is a ubiquitous component of indoor and outdoor air pollution that is epidemiologically linked to many human pulmonary diseases. PM has many emission sources, making it challenging to understand the biological effects of exposure due to the high variance in chemical composition. However, the effects of compositionally unique particulate matter mixtures on cells have not been analyzed using both biophysical and biomolecular approaches. Here, we show that in a human bronchial epithelial cell model (BEAS-2B), exposure to three chemically distinct PM mixtures drives unique cell viability patterns, transcriptional remodeling, and the emergence of distinct morphological subtypes. Specifically, PM mixtures modulate cell viability and DNA damage responses and induce the remodeling of gene expression associated with cell morphology, extracellular matrix organization and structure, and cellular motility. Profiling cellular responses showed that cell morphologies change in a PM composition-dependent manner. Lastly, we observed that particulate matter mixtures with high contents of heavy metals, such as cadmium and lead, induced larger drops in viability, increased DNA damage, and drove a redistribution among morphological subtypes. Our results demonstrate that quantitative measurement of cellular morphology provides a robust approach to gauge the effects of environmental stressors on biological systems and determine cellular susceptibilities to pollution.

## Introduction

Ambient air pollution threatens human health through direct links to chronic illnesses and premature deaths. High pollution levels are associated with elevated incidences of ischemic heart disease, lung cancer, aggravated asthma, chronic obstructive pulmonary disease (COPD), stroke, and adverse birth outcomes(1–6). In 2019, it was estimated that 6.67 million deaths could be attributed to air pollution exposure worldwide(7). Particulate matter (PM), which consists of microscopic solids and liquid droplets, is an important component of ambient air pollution. These particulates and their precursor chemicals are emitted from many natural and man-made sources, including volcanic activity, burning of biomass, vehicle emissions, coal-burning powerplants, and other industrial activities(8). However, the molecular mechanisms governing how cells change in response to air pollution remain poorly understood.

Recent studies have identified strong associations between PM size and different biological responses(9, 10). However, a key challenge in elucidating the health effects of PM exposure is that PM chemical composition can vary greatly across geographical areas and environments, as there are various anthropogenic and biogenic contributors that emit different chemical species(11–13). These inherent geographical differences of PM can impose challenges towards understanding the different influences at the cellular and molecular levels, since the biological effects can vary with chemical composition. Studies using lung cell models such as A549, BEAS-2B, or primary airway epithelial cells have demonstrated the impact that different PM mixtures and pollutants can have on cellular pathway remodeling(10, 14). For example, in parallel studies a bronchial epithelial cell model, BEAS-2B, exposed to mixtures of either secondary organic aerosol or aerosolized formaldehyde showed unique molecular responses and pathway remodeling(15, 16). Additionally, other studies have investigated the induction of oxidative stress due to particulate matter exposure and have highlighted unique regulatory pathways that contribute to the pro-inflammatory response(14, 17).

Different exposure methods have also investigated the biological effects of PM including liquid submerged exposures(18), air-liquid-interface exposures (ALI)(19, 20), and pseudo-air-liquid-interface exposures(18). It is worth noting that these vary in cost, physiological relevance, and throughput. Studies have also looked at a variety of environmental pollutants including PM_10_, PM_2.5_, and PM_0.1_ (particulate matter with aerodynamic diameters of less than 10µm, 2.5µm, and 0.1µm, respectively) collected from cities including Beijing, Milan, Seoul and others(21–26). Organic and aqueous extractions of PM have also been investigated along with individual components or pollution types including secondary organic aerosols, diesel exhaust particles, volcanic ash, and metals. However, the results of these studies vary greatly, in part, due to their use of different cell models, exposure times and protocols, and PM types that are often not fully characterized. All these factors introduce challenges to drawing meaningful comparisons of the biological effects of different PM types.

Previous studies looking at air-pollution-induced pathway remodeling via transcriptomics have found changes in regulatory pathways that control cellular morphology, including significant alterations in cholesterol synthesis pathways of bronchial epithelial cells that result in distinct morphological changes(16, 27). By extension, these types of studies indicate that a retraction in cell size could be used as a biomarker of toxicity(16). Overall, cellular and nuclear morphology is linked to upstream changes in gene expression and cellular dysfunction(28, 29), with significant pathway remodeling in cell death programs, apoptotic pathways, extracellular matrix (ECM) interactions, and cytoskeleton structures(29–31). In the context of aging, increases in cell and nuclear sizes, as well as irregularities in cell shapes associate strongly with fundamental defects and senescence(32, 33). While longer term pollutant exposures of lung cells are linked to increased senescence, it is unclear how short-term exposures modulate cellular responses based on molecular or morphological phenotypes(34).

Here, we expose the BEAS-2B human bronchial epithelial cell model to three well-characterized and compositionally unique PM mixtures available from the National Institute of Standards and Technology (NIST): Urban (SRM1648a), Fine (SRM 2786) and Diesel Exhaust (SRM 2975). Exposures were performed at multiple concentrations ranging from 31 to 1000 µg/mL for 24 hours to investigate the effects of multiple PM types on human lung epithelial cells. Following exposures, we measured transcriptional changes to identify specific PM-composition-dependent remodeling of molecular pathways. In parallel, we performed morphological analysis of cells at baseline and after PM exposures to develop a robust single-cell platform to profile cellular responses and the emergence of functional subtypes of cells. Together our study provides a multi-scale approach to quantify molecular and morphological responses to several relevant PM mixtures. Additionally, we show that cell morphology can encode susceptibility to particulate matter exposure, offering a new tool for understanding the cellular effects of environmental stressors.

## Results

### Cellular viability is differentially affected by unique particulate matter samples

To determine the biological effects of different PM compositions on cell viability, we measured the survival of BEAS-2B cells following exposure to three individual PM mixtures sourced from NIST (Urban (SRM1648a), Fine (SRM 2786), and Diesel Exhaust (SRM 2975)) to quantify changes in toxicity to cells. The Urban and Fine samples contain PM collected over extended periods of time from two different cities, St. Louis, Missouri and Prague, Czech Republic, respectively. The Diesel Exhaust sample was collected from the exhaust of a diesel-powered engine. Importantly, these mixtures exhibit major differences in several components; for instance, the mass fractions of lead, cadmium, and nitro-polycyclic aromatic hydrocarbons (nitro-PAHs) vary by at least an order of magnitude between at least two of the samples (Table 1). A complete comparison of reported compositional data can be found in Dataset S1. Interestingly, cadmium and lead are both highly toxic metals that can be found in air pollution from manufacturing of batteries, cigarette smoke, metal processing, and production of plastics(8, 35), while nitro-PAHs are primarily emitted from combustion of diesel fuel and have been shown to have mutagenic and genotoxic properties(36). Diesel exhaust is a major component of air pollution in urban areas resulting from the heavy traffic, and diesel engines emit more particles and 10-times higher levels of nitro-PAHs than gasoline engines(37).

**Table 1.** Select compositional differences between PM types, as reported by NIST in mg per kg of total particulate matter mass. No cadmium or lead concentrations were reported for SRM 2975.

| Units:<br>mg/kg | Urban<br>PM mass<br>fraction | Fine PM<br>mass<br>fraction | Diesel<br>Exhaust<br>mass<br>fraction |
| --- | --- | --- | --- |
| Cadmium | 73.7 | 4.34 | - |
| Lead | 6550 | 286 | - |
| Nitro-PAHs | 0.73962 | 0.99598 | 45.907 |

To evaluate the effects of PM exposures on cellular viability, we used the alamarBlue™ viability assay. We observed that after cells were exposed to PM for 24 hours (Fig. 1a), cell populations exhibited PM-type- and concentration-dependent changes in viability (Fig. 1b). For example, Urban PM induced a steady decrease in viability at concentrations greater than or equal to 250µg/mL (p<0.05). However, Fine PM induced a significant decrease in viability only at the highest concentration of 1000 µg/mL. Paradoxically, diesel exhaust PM induced an increase in cell viability across all concentrations. The exposure concentrations of 125 and 500 µg/mL, equivalent to 35.2 µg/cm^2^ and 140.8µg/cm^2^ in terms of deposition over cell growth area, were chosen for further analysis. These concentrations were chosen based on previous analyses indicating that 20 µg/cm^2^ could be deposited in the tracheobronchial regions of the lung over a period of 8 hours in an urban environment(38), ∼35.2 µg/cm^2^ falls within an expected deposition amount within areas of the human lung for a 24-hour period in an urban environment, and ∼140.8µg/cm^2^ could be representative of exposure levels in extremely polluted cities.

**Figure 1.**
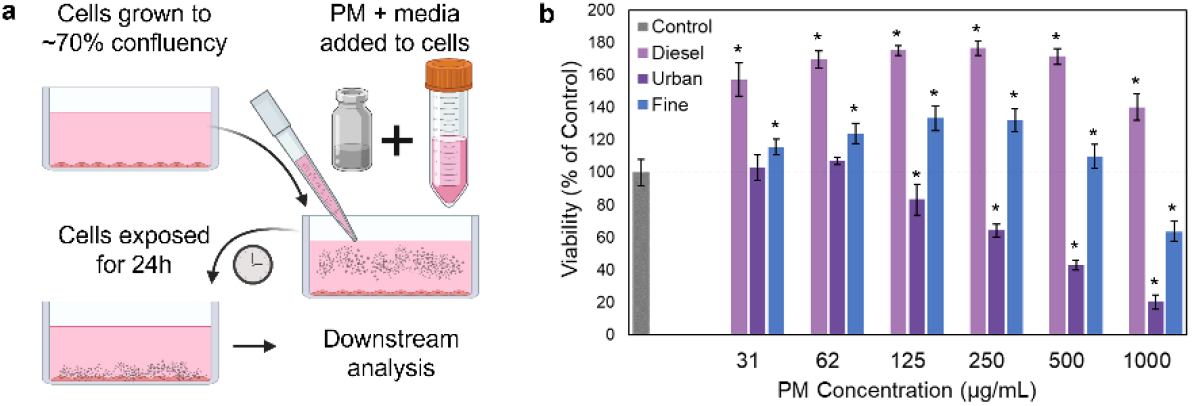
Effects of particulate matter (PM) exposure on cell viability. (**a**) Graphical depiction of submerged PM exposure method. (**b**) Cell viability following 24h exposures to different PM types and concentrations. Values are percentages of viable cells relative to unexposed control cells as measured with the alamarBlue assay (n=7, error bars represent one standard deviation, * = p<0.05 using Student’s T-test).

### Exposure to different PM types and concentrations induce differential DNA damage responses and cell death

Interestingly, the alamarBlue™ viability assay did not show decreases in viability for Fine and Diesel PM exposures across a wide range of concentrations (up to 500 µg/mL). To investigate this further, we profiled the DNA damage responses to PM mixtures. Using confocal microscopy, we measured the accumulation of the histone phosphorylation γH2AX as a marker of double stranded DNA breaks, a precursor to genotoxicity and cell death(39). We found that exposures to the different PM mixtures at concentrations of 125 or 500 µg/mL led to differential levels of DNA damage, with all exposures leading to increases in DNA damage relative to the control, unexposed cells (Fig. 2a,b).

**Figure 2.**
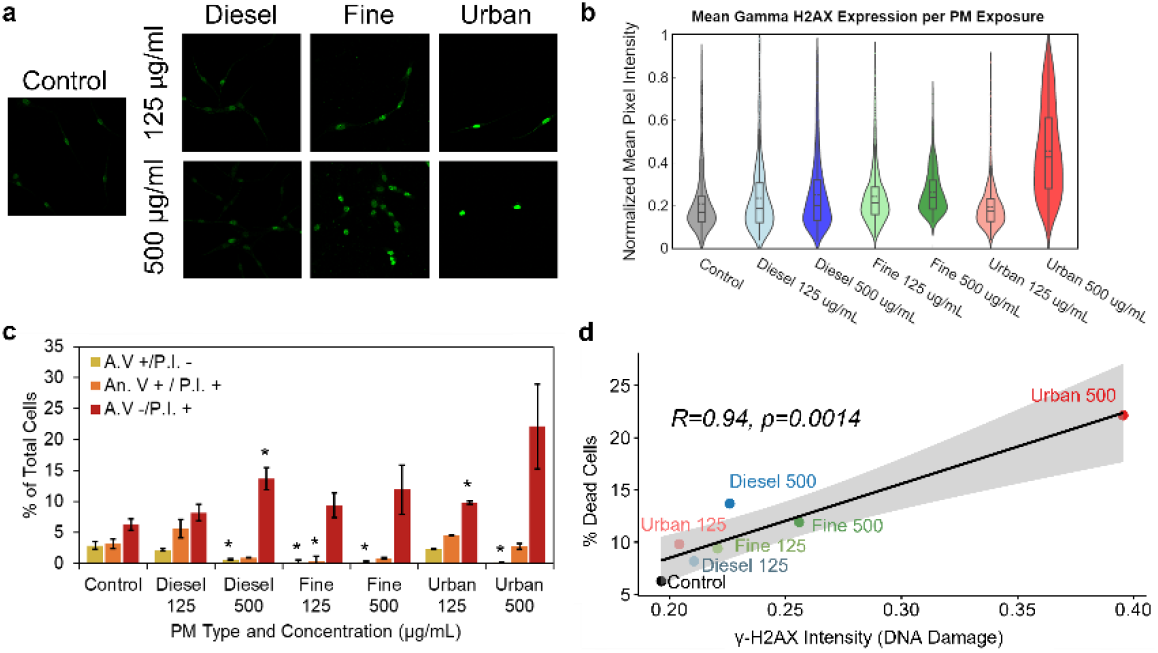
PM exposure leads to alteration of apoptotic levels and DNA damage. (**a**) Representative images of the immunofluorescent staining of γH2AX across different exposure conditions. (**b**) Violin plots overlaid with box and whisker plots showing the distribution of average γH2AX intensity values for cell nuclei in each exposure condition. (**c**) Flow cytometry analysis of Annexin V-Propidium Iodide apoptosis assay following PM exposure. A.V+/P.I.- and A.V+/P.I.+ represent early- and late-stage apoptotic cells respectively, A.V-/P.I.+ represents dead cells (n=3, ≥10,000 cells per measurement, error bars represent the standard error of the mean, * = p<0.05). The remaining cells in each condition were healthy (A.V-/P.I.-). (**d**) Correlation between percentage of dead cells from the apoptosis assay shown in (**a**) and the average γH2AX intensity following PM exposure. Gray shading represents a 95% confidence interval.

Furthermore, we evaluated whether PM exposures were inducing cell death via apoptosis or based on non-apoptotic mechanisms. Both apoptotic and non-apoptotic mechanisms are associated with aberrant levels of DNA damage. To determine the mode of cell death, we incubated cells with Annexin V (A.V) and Propidium Iodide (P.I.) after exposure to the different PM mixtures and quantified the levels of apoptotic and dead cells using flow cytometry, as previously used to investigate the mode of cell death in lung cells exposed to PM mixtures(40).

Cells exposed to 125 µg/mL of all PM conditions exhibited small increases in the population of dead cells (A.V-/P.I.+) and a decrease in the population of apoptotic cells (A.V+/P.I.- and A.V+/P.I.+) (Fig. 2c). A.V+/P.I.-indicated cells were in the early stages of apoptosis, while A.V+/P.I.+ indicate cells are in later stages of apoptosis or dying due to loss of membrane integrity. In contrast, exposure to 500 µg/mL concentration of each PM mixture resulted in an increase in the number of dead cells (A.V-/P.I.+) (Fig. 2c). Representative scatter plots of flow cytometry data from each condition are shown in Fig. S1. We next compared the trends in the populations of apoptotic versus dead cells across the different PM mixtures and observed that exposure to different PM compositions led to different distributions of apoptotic versus dead cells. For instance, exposure to the Urban PM mixtures resulted in a greater number of dead cells relative to the Fine and Diesel mixtures. These results corroborate the general pattern in viability observed via the alamarBlue™ assay, with Urban PM inducing the greatest losses of viability, but better captures loss of viability in the Diesel and Fine conditions (Fig. 1b).

The levels of DNA damage are also associated with the levels of observed cell death (Fig. 2d). As shown in Figure 2, γH2AX intensity increases with exposure to increasing PM concentrations for each of the three PM types. Furthermore, γH2AX intensity positively correlates (Pearson Coefficient of R=0.94, p=0.0014) with the percentage of dead cells (A.V-/P.I.+) found in the Annexin-V Propidium iodide data for the same exposed populations (Fig. 2d). Taken together, these data show increases in cell death and DNA damage levels are observed with increasing PM concentrations. These levels are also dependent on the PM composition, as the three mixtures show markedly different trends. Additionally, the correlation between cell death and γH2AX intensity points to a framework of DNA damage associated cell death.

### Post-exposure transcriptional remodeling of cell populations in response to PM mixtures indicates common and unique gene expression activation

The unique differences observed in viability and in the patterns of DNA damage following PM exposures at 125 and 500 µg/mL prompted us to investigate whether PM exposures also induced differential molecular responses. To better understand how the underlying transcriptomic profiles influence differential viability across PM mixtures, we assessed changes in gene expression patterns via 3’-TagSeq(41, 42) (Datasets S2-7). This approach takes advantage of the poly(A) tail on mRNA for sequencing library preparation, allowing the accurate quantification of protein coding transcripts.

We first observed that exposure to Urban and Fine PM mixtures induced significant changes in the expression of a greater number of mRNA transcripts relative to Diesel Exhaust under the two PM concentrations tested. Furthermore, the magnitudes of the changes were larger for cells exposed to Urban and Fine mixtures than those exposed to Diesel Exhaust PM (Fig. 3a-f). These observations indicate that the Urban and Fine mixtures have a lower threshold for stimulation of cellular responses. Additionally, the number of genes that were differentially expressed by each PM type increased with higher concentrations (*i.e.*, 125 µg/mL versus 500 µg/mL exposures) (Fig. 3a-f). Moreover, the majority of the genes (at least 67% for each condition) that were up- and down-regulated in the 125 µg/mL conditions were similarly up- and down-regulated in the 500 µg/mL condition (Fig. S2), indicating consistency in the transcriptional responses across different concentrations of each PM type, with additional pathway activation at higher concentrations.

**Figure 3.**
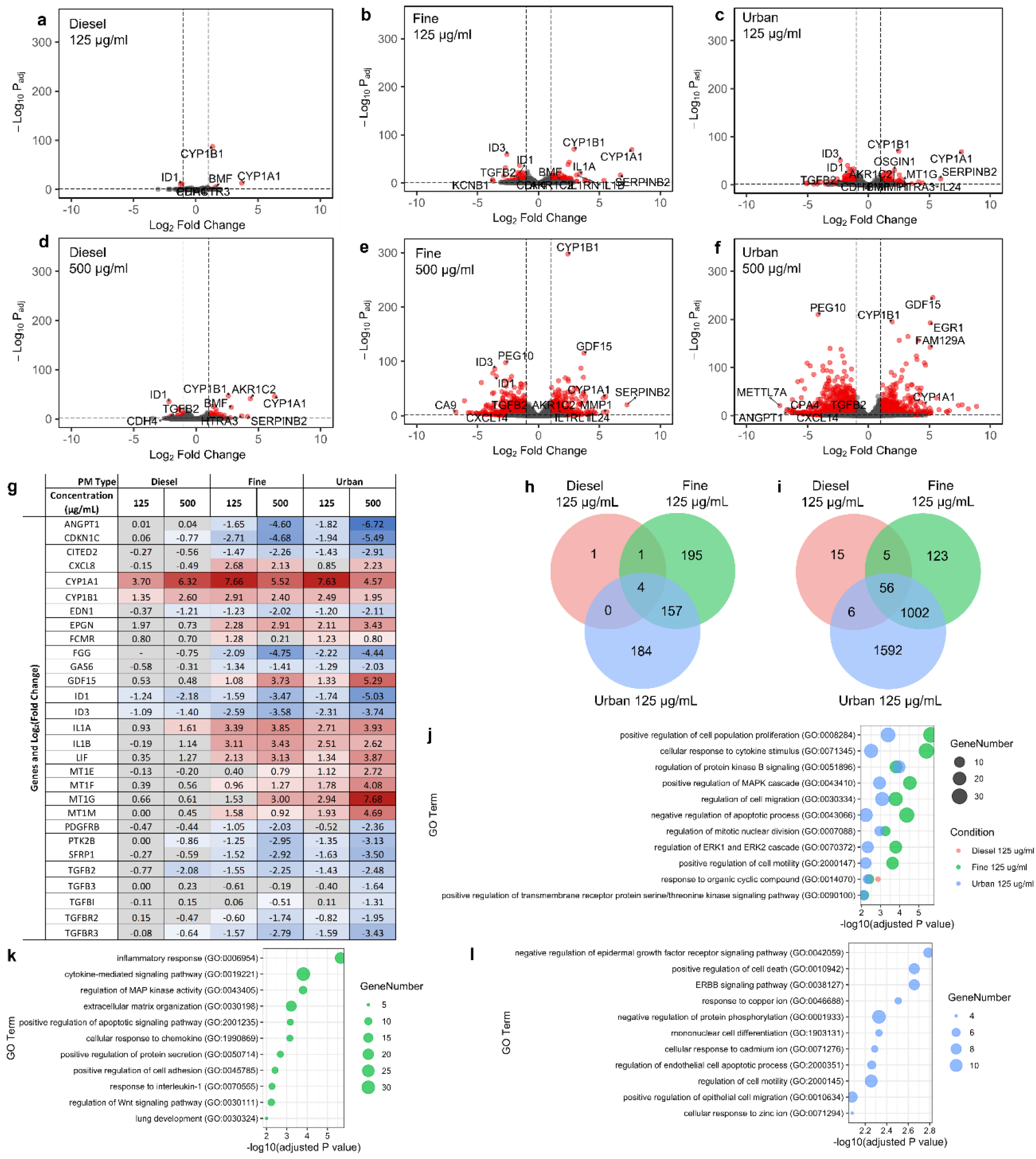
Transcriptomic analysis of particulate matter stress reveals unique network remodeling. (**a-f**) Volcano plots showing significantly differentially expressed (DE) genes (red = Log_2_FC>1, p_adj_<0.05) relative to control cells after exposure to Diesel, Fine and Urban PM at 125µg/mL (**a-c**) and 500µg/mL (**d-f**) for 24 hours. (**g**) Log_2_Fold Changes in expression of select genes. A grey background indicates the expression change was not significant (p>0.05). (**h**, **i**) Venn diagrams of the significantly DE genes from each condition. Intersections represent genes that were differentially expressed in overlapping conditions. (**j-l**) Bubble plots showing select enriched Gene Ontology (GO) Biological Process terms that are commonly enriched among two or more of the low-level exposure conditions (**j**), or unique to the low-level Fine **(k)** or Urban **(l)** exposure conditions.

Additionally, we observed that four mRNAs encoded by the CYP1A1, CYP1B1, ID1, and ID3 genes were differentially expressed post-exposure across all conditions (Fig. 3g); two were overexpressed (CYP1A1 and CYP1B1) and two displayed decreased expression (ID1 and ID3), relative to expression levels in unexposed control cells. The CYP1A1 and CYP1B1 are members of the cytochrome P450 family that are involved in the metabolism of endogenous compounds such as fatty acids and steroid hormones(43). Consistent with our results, these genes are upregulated in human epithelial lung cell models in response to exogenous polyaromatic hydrocarbons (PAHs) present in PM(14). These PAHs bind to the cytosolic aryl hydrocarbon receptor (AhR), which then mediates expression of the cytochromes and promotes a proinflammatory response to induce ROS production in cells. ID1 and ID3 are inhibitors of DNA binding proteins that are induced by TGF-β and have been implicated in regulation of senescence, apoptosis, and cell cycle alterations(44). Moreover, ID1 expression has also been shown to decrease after exposure to coarse PM (PM with an aerodynamic diameter between 2.5-10 µm)(10), but the roles of these genes have been less defined in the context of air pollution exposures. Importantly, the increase in expression of the CYP genes and the decrease in expression of the ID genes suggest that the response to organic cyclic compounds as well as alteration of the TGF-β regulatory pathways are commonly remodeled by these unique PM mixtures. This is further supported by the differential expression of additional TGF-related genes (Fig. 3g). Interestingly, many TGF-β related genes are involved in the regulation of cell morphology and motility.

We next observed that unique mRNAs were significantly differentially expressed only when cells were exposed to certain PM mixtures, but not others (Fig. 3h,i). For example, TNFAIP6, a regulator of the ECM, LCAT, a protein involved in extracellular metabolism, and CXCL1, a protein involved in inflammation, are significantly differentially expressed under only the Fine exposure conditions. However, genes including DDIT4, a protein induced by DNA damage, MT1E, a protein involved in the cellular response to cadmium, and ACTN4, an actin binding protein, are differentially expressed under only the Urban exposure conditions. This indicates that there could be unique pathway activation that is dependent upon the PM composition.

Overall, the gene expression patterns observed in cells exposed to Fine and Urban PM mixtures exhibit significant pathway remodeling, whereas cells exposed to Diesel Exhaust PM exhibit less remodeling. Similarly, we observed a dose-dependence in the extent of pathway remodeling, i.e., more changes with higher PM concentrations. Lastly, we noted that, although expression of a limited set of four genes was consistent across all conditions (*i.e.*, CYP1A1, CYP1B1, ID, and ID3), other genes are differentially expressed in a manner that is dependent on the PM type.

### Gene Ontology analysis reveals PM-dependent remodeling of apoptosis, motility, and morphology pathways

To determine the key remodeled pathways post-exposure and the extent to which they were remodeled, we performed Gene Ontology (GO) and pathway enrichment analysis. We performed this analysis using the transcriptomics data from cells exposed to Urban, Fine, and Diesel Exhaust PM mixtures at 125 and 500 µg/mL (Fig. 3j-l, Fig. S3). The complete list of enriched GO Terms for each condition can be found in Datasets S8-S13. Using Enrichr(45), we identified 34 pathways that were significantly enriched (p_adj_<0.01) in cells exposed to the 125 µg/mL concentration of both Urban and Fine PM mixtures, relative to baseline. We selected 11 non-redundant pathways to show in Fig. 3j. We observed changes in the expression of genes related to MAPK cascade (*e.g.*, EDN1, GDF15, TGFB2, ANGPT1, and LIF), epithelial cell proliferation, (*e.g.*, CDKN1C and EPGN), regulation of apoptosis (*e.g.*, FCMR and CITED2), and cell migration and extracellular matrix organization pathways (*e.g*., SFRP1 and FGG). It is worth noting that for cells exposed to Urban and Fine PM at 500 µg/mL, similar pathways were also significantly enriched (p_adj_ <0.01) (Fig. S3, Datasets S11-S13).

Interestingly, we observed few differentially expressed genes in cells exposed to the Diesel Exhaust PM mixture. Only the “response to organic cyclic compound” pathway was significantly enriched (p_adj_ <0.01) in cells exposed to all PM types at the 125 µg/mL concentration. However, this response appears to be ubiquitous, with the CYP1A1 and CYP1B1 genes increasing in expression across all conditions post-exposure. Similarly, we identified upregulation of IL1B, which was upregulated in all conditions except 125 µg/ml Diesel Exhaust PM.

We also identified key genes exhibiting differential expression across both the 125 µg/mL Urban and Fine PM exposure conditions that contributed to the remodeling of multiple pathways (Figs. 3i-j, S4). For example, IL1A, IL1B, and TGFB genes were part of several Gene Ontology-defined pathways that comprise cytokine signaling cascade and TGF-β signaling. Other genes involved across many pathways include GAS6, which is involved in cell growth and migration and cytokine signaling, and PTK2B, a protein involved in the activation of MAPK signaling and reorganization of the actin cytoskeleton. These genes are present in many of the most significantly altered pathways, highlighting their importance in the biological response to PM exposure.

Lastly, we observed that several exclusive GO terms were significantly enriched (p_adj_<0.01) in cells post-exposure to Urban PM at both concentrations (125 and 500 µg/mL) (Fig. 3k, Fig. S3) that included unique responses to metal ions. Examples of these pathways include response to cadmium ion, copper ion, and zinc ion, which encompass mRNAs encoded by the MT1 family genes (MT1G, MT1E, MT1F, and MT1M). The patterns of gene expression changes involved in the regulation of metal ions are consistent with the increase in metal composition (*i.e.*, cadmium) in the Urban PM mixture, relative to the other mixtures tested (Table 1). Similar to the 125 µg/mL Urban exposure, at the 500 µg/mL Urban condition, the top significantly enriched GO term is response to metal ion, again indicating the importance of the increased metal concentrations in the Urban PM sample relative to Fine and Diesel Exhaust.

Taken together, these data indicate that cells differentially regulate their gene expression patterns in a PM composition dependent manner. However, pathways related to cell morphology, and extracellular matrix remodeling seem to be broadly shared across all PM exposure conditions, with pathways related to apoptosis shared across the Urban and Fine conditions.

### Particulate matter compositions drive the emergence of morphological subtypes post-exposure

Since unique PM mixtures drive differential responses, particularly in apoptosis, cytoskeletal structure, and ECM-related pathways, we wondered whether these responses could be captured by changes in cellular morphologies across cell populations. Using our BEAS-2B cell line model, we exposed cells to the same PM mixtures at the same concentrations and exposure times. After exposure, cells were fixed and stained for F-Actin (488-Phalloidin), DNA (DAPI), and γH2AX (anti-γH2AX (phospho-S139) antibody) (Fig. 4a,b). The Phalloidin and DAPI stains were used to delineate the cell and nuclear boundaries, and γH2AX to quantify the extent of persistent DNA damage. For each cell and nuclear boundary, we computed 33 discrete parameters describing features related to the sizes and shapes of individual cells (Table S1). Across all conditions we analyzed ∼13,000 single cells. To identify whether BEAS-2B cells exhibited morphological subtypes that changed after PM exposure, we performed dimensional reduction and clustering analyses on cells analyzed across all conditions. Using a combination of k-means clustering and Uniform Manifold Approximation and Projection (UMAP), we identified 10 distinct morphology clusters, each having unique cellular and nuclear morphological profiles (Fig. 4c,d). Furthermore, these ten morphological clusters can be further grouped into three cluster groups (CG), with morphology clusters 1, 2, and 5 belonging to CG1, morphology clusters 3, 4, and 6, belonging to CG2, and morphology clusters 7-10 belonging to CG3 (Fig. S5).

**Figure 4.**
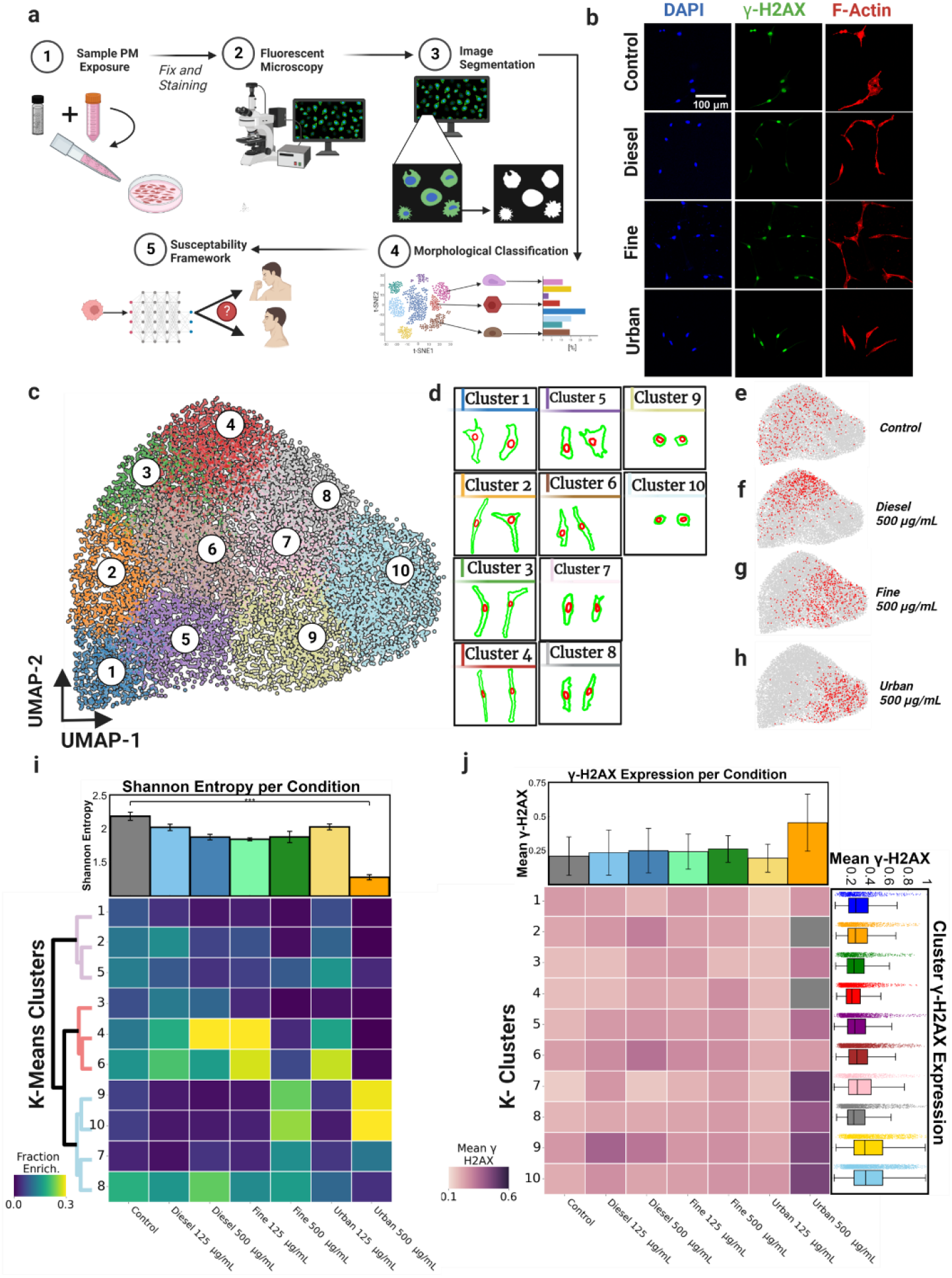
(**a**) Graphical depiction of morphological analysis pipeline. (**b**) Representative fluorescence microscopy images of cells from each condition. (**c**) UMAP visualization of the 33 measured morphological parameters for each cell in every condition. UMAP-1 (X-axis) was *negatively* correlated with size and UMAP-2 (Y-axis) was *positively* correlated with cell elongation, or linearity. k-means clustering was applied to cluster cells of similar morphologies. (**d**) Representative cellular (green) and nuclear (red) morphologies of cells from each k-means morphology cluster. (**e-h**) Plots showing the distribution of cells from each respective exposure condition in red within the UMAP space. (**i**) Heatmap displaying the enrichment in number of cells in each morphology cluster for each exposure condition. The bar graph shows the Shannon Entropy for the distribution of cell morphologies within each exposure group. The dendrogram identifies clusters with similar morphological features. (**f**) Heatmap displaying the mean γH2AX in morphology clusters across all exposure conditions. Mean γH2AX intensity across each exposure condition (top). γH2AX intensity of all cells within each k-means cluster (right).

Next, we asked whether cells exposed to both low (125 µg/mL) and high (500 µg/mL) concentrations of each PM mixture exhibited differential abundance of cells across each morphology cluster. Upon comparison, we observed pronounced shifts in the abundance of cells per morphology cluster in a PM-dependent manner (Figs. 4e-i and S6). Specifically, when compared with unexposed conditions, cells exposed to 500 µg/mL of Urban and Fine PM exhibited higher fractions of cells in clusters 9 and 10, which describe smaller, more rounded morphologies (Fig. 4g-i). However, cells exposed to 500 µg/mL of Diesel PM exhibited higher fractions of cells in clusters 4 and 8, which describe larger, more elongated cell morphologies (Fig. 4f,i). Based on the observed fractional redistributions among morphology clusters per condition, we computed the Shannon entropy as a way to estimate cellular heterogeneity(46). Although cells redistributed among morphology clusters per PM conditions, only cells exposed to Urban 500 µg/mL showed a pronounced decrease in heterogeneity relative to unexposed control cells (Fig. 4i).

Taken together, our results indicate that cells exposed to different PM mixtures drive fractional redistributions among cellular morphology clusters in a PM-dependent manner. Furthermore, the differential localization of cells exposed to Urban PM (small, more rounded morphologies) and Diesel PM (larger, more elongated morphologies) point out that these PM mixtures are likely driving unique responses based on the underlying compositions. Lastly, these results suggest the potential utility of cell morphology cluster profiles to denote functional subtypes in pre- and post-exposed cells.

### Morphological clusters are further defined based on the extent of persistent DNA damage

Given that cells exposed to both Urban and Fine PM exhibited a higher fraction of cells with smaller, more rounded cell morphologies (Fig. 4g,h) and decreased viability relative to unexposed cells (Fig. 1b), we investigated whether morphology clusters were associated with persistent DNA damage. Since each cell was co-stained for γH2AX, we computed the extent of DNA damage based on the total nuclear abundance of phosphorylated-H2AX (γH2AX). Comparing cells from all exposure conditions, we observed a significant increase in the γH2AX content for cells exposed to 500 µg/mL Urban PM relative to unexposed control cells. Furthermore, to test whether cells in different morphology clusters exhibited different levels of DNA damage, we pooled cells within each morphology cluster across all conditions and quantified the levels of γH2AX. Interestingly, we found that cells belonging to clusters 9 and 10 had the highest levels of γH2AX (*i.e.*, high DNA damage), with cluster 4 exhibiting the lowest level of damage (Fig. 4j). These results suggest that the identified cell morphology clusters could be further defined based on the extent of DNA damage and susceptibility to cell death after PM exposure.

### Cadmium drives morphological shifts among functional clusters after PM exposures

To further test the hypothesis that the chemical compositions of the PM mixtures drive specific shifts among morphological clusters (*i.e*., smaller, rounder, and less viable cells), we systematically supplemented our PM mixtures with different concentrations of cadmium chloride (0-25µM) and lead acetate (0-250µM) that mimic those used in other studies(47, 48). We selected cadmium (Cd) and lead (Pb), due to their variable concentrations across different PM mixtures (Table 1), and the pronounced shifts in both the viability and the morphological shifts when cells were treated with Urban PM (Urban PM has the highest concentration of Cd in the tested PM mixtures). First, we observed a significant decrease (p≤0.05) in viability with increasing levels of cadmium chloride supplementation across all conditions tested (Figs. 5a, S7). In contrast, cells exposed to PM mixtures supplemented with lead acetate resulted in little to no change in viability (Fig. S8).

**Figure 5.**
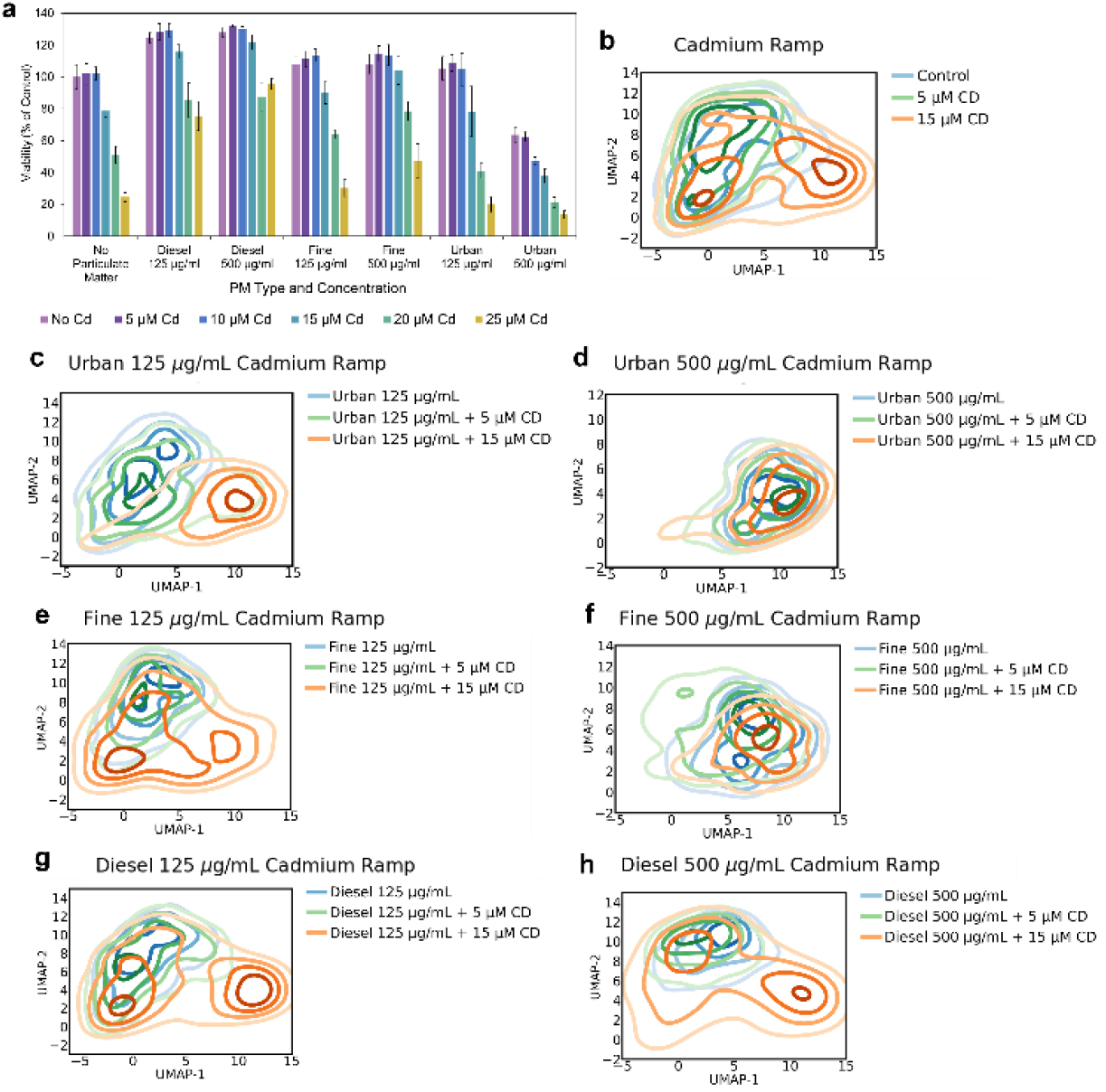
(**a**) Cell viability following 24h exposures to different PM types and concentrations supplemented with cadmium chloride (0-25 µM Cd). Values are percentages of viable cells relative to unexposed control cells as measured with the alamarBlue assay (n=6, error bars represent one standard deviation). (**b-h**) Morphological distribution of cells from each respective exposure condition with CdCl_2_ supplementation displayed across the UMAP space.

Evaluating the morphological effects of cells exposed to increasing CdCl_2_ concentrations across all PM mixtures (Fig. 5b-h), we observed a general tendency towards smaller, rounded morphologies described by clusters 9 and 10. Cells exposed to 125 µg/mL Urban PM and 15µM Cd resembled the distributions of 500 µg/mL Urban either alone or with 5 or 15µM cadmium supplementation (Figs. 5c,d and S9b,c). For cells exposed to 125 µg/mL of Fine PM, 15µM cadmium supplementation led to a great shift relative to the 5µM. However, in the 500 µg/mL Fine PM conditions, even at 5µM we observed a shift towards clusters 9 and 10, with 500 µg/mL Fine PM with cadmium supplementation resembling the 500 µg/mL Urban PM conditions (Figs. 5e,f and S9d,e). Lastly, cells exposed to 125 µg/mL Diesel PM with 15µM cadmium exhibited a bi-phasic shift in the abundance of cells among clusters, with 33.9% of cells shifting towards clusters 9 and 10. However, cells exposed to 500 µg/mL of Diesel and 15µM cadmium exhibited a similar shift towards clusters 9 and 10 (30.4% of cells), despite the increased PM concentration (Fig 5g,h).

Collectively, our data indicates that the differential abundance of cadmium in the different PM mixtures may drive differential toxicity among PM mixtures. Importantly, these observed correlations between increased PM toxicity (lower viabilities with cadmium supplementation) and distinct morphological redistributions among cell populations suggest the potential for predicting the toxicity and susceptibility of cells to different PM mixtures using their morphologies.

### Single cell morphology predicts susceptibility to Urban PM exposures

Since cells exhibited unique morphological phenotypes and responses to PM exposures, we wondered whether cellular morphologies encoded resilience or reduced susceptibility to PM exposure at the single-cell level. To test whether the starting morphologies of cells associated with the response to PM exposure, we isolated single-cell clones from the parental BEAS-2B cell line. Seeding a single cell per well of a 96 well plate, we generated twelve single-cell clones. Analyzing the morphologies of each clone, we did not observe any clone localizing specifically to one morphology cluster (Fig. 4a). However, when separating the individual morphological clusters into the three cluster groups (CG1, CG2, CG3), we observed that some clones occupied primarily one or multiple of the three cluster groups (Figs. 6a-c, S10a, and S11). As expected, when we compared the cellular heterogeneities of the twelve clones relative to the parental, we observed an overall reduction in the overall Shannon entropy for each of the clones, with clones 7 and 1 having the lowest heterogeneity (Fig. S12).

**Figure 6.**
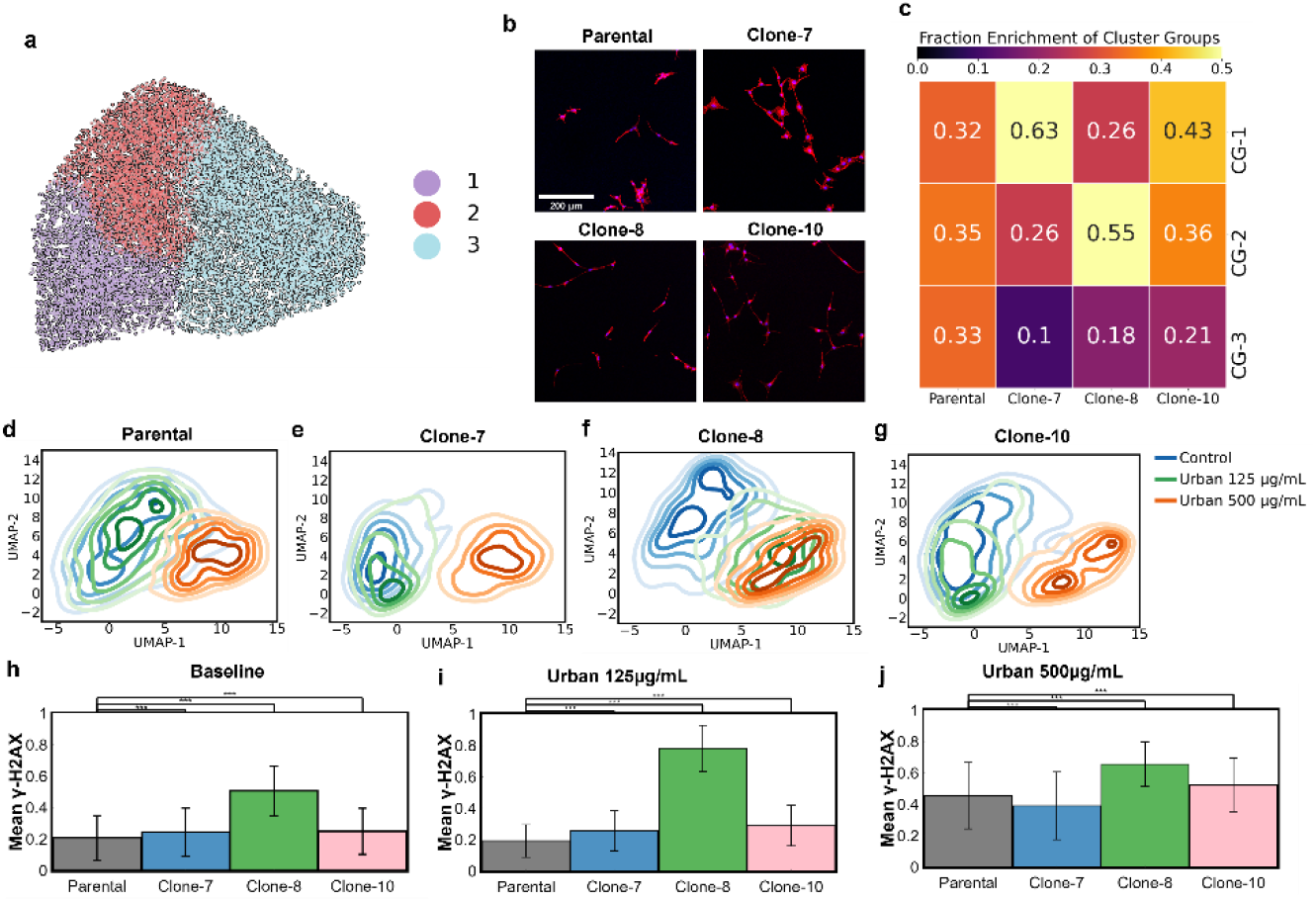
Morphology encodes susceptibility to particulate matter exposure. (**a**,**b**) The 10 k-means clusters that are used to define cell morphology can be further grouped into 3 cluster groups (CG1-3) using hierarchical clustering. (**c**) Clonal populations show enrichment in different morphological cluster groups. The distributions of cell morphologies differ for each clone and the parent cell population from which the clones were derived. (**d**-**g**) Upon exposure to Urban PM, clones with unique baseline morphologies show different shifts in morphology. (**h**-**j**) Average γH2AX intensity values in cell nuclei for populations in each exposure condition.

To further test the hypothesis that the starting cellular morphologies governed the response to PM mixtures we exposed all clones to the Urban PM mixture at 125 and 500 µg/mL for 24 hours. To illustrate unique baseline morphologies, we selected the parental and three clones that exhibited differential abundance of cells within the three morphological cluster groups. Specifically, at baseline the parental line had a similar abundance of cells across all three cluster groups, clone-7 was highly abundant for cells in CG1, clone-8 was highly abundant in CG2, and clone-11 was highly abundant in both CG1 and CG2 (Fig. 6c). Based on these starting morphologies we tested the cellular responses to Urban PM exposures (both 125 and 500 µg/mL).

After exposure, single cell clones showed differences in morphological distributions. For the parental and isolated clonal populations, cells exposed to 500 µg/mL of Urban PM resulted in a drastic shift towards CG3 and more specifically morphology clusters 9 and 10. However, the major differences as a function of baseline morphologies were observed in the cells exposed to 125 µg/mL of Urban PM (Fig. 6d-g, Fig. S10b). For the parental, clone-7 and clone-11 populations, there were very little shifts in the abundance of cells within the cluster groups, as shown by the significant overlap of contours from control (unexposed) and the 125 µg/mL conditions (Fig. 6d,e,g). Interestingly, for clone-8, at 125 µg/mL Urban PM there was a significant shift towards CG3 (specifically clusters 9 and 10). These results suggest that c8 may be more susceptible to Urban PM relative to clone-7 and clone-11 (Fig. 6f). To further test this notion of susceptibility, we evaluated whether DNA damage responses contributed to the susceptibility. For cells in the parental, clone-7, and clone-11 populations there was low expression of γH2AX which increased significantly at 500 µg/mL of Urban PM, indicating increasing DNA damage relative to baseline. However, clone-8 exhibited high γH2AX signal at baseline (>2.4x compared to the parental, p<0.001), with a large increase at both concentrations of Urban PM (p<0.001) (Fig. 6 h-j).

Taken together, these results point to the notion that baseline morphologies encode susceptibility to Urban PM exposure, and generally that baseline cell morphology profiles can be used as predictors/biomarkers of PM-induced responses. Clones with a high abundance of cells in CG1 were most resilient to Urban PM, while clones having a high abundance of cells in CG2 and CG3 demonstrated increasing susceptibility to Urban PM. Lastly, susceptibility to Urban PM exposures seem to be influenced by the baseline levels of DNA damage.

## Discussion

In this study, we demonstrate a multi-scale approach to characterize the unique differences in cellular response to three PM mixtures using molecular and quantitative morphological analyses. We further investigated morphological variations across populations of unexposed and PM-exposed cells to show that cellular morphology encodes susceptibility to Urban PM exposures and provides mechanistic insights into variable responses across cell populations. Additionally, we show that these responses are dependent on the composition of the PM mixture, for instance, abundance of cadmium can drive unique cellular transcriptional responses and morphological changes.

With the emergence of single-cell technologies and deep-learning tools, there has been a tremendous acceleration in the capacity to quantify and analyze specific cell states and behaviors across cell populations(49–51). Specifically, analysis of biophysical properties, such as motility and morphology, offer an efficient method to discretize functional subtypes of cells(32, 52, 53). In this work we profile the single-cell morphological changes after exposure to various PM mixtures to quantify cellular responses and identify cellular properties that are associated with cellular susceptibility to pollutants. Three major and novel findings of this work include the following: First, we identified that although there is a common transcriptomic response to PM in the activation of the P450 family cytochromes, as shown previously(14, 54), the degree of pathway remodeling is dependent on the PM composition and concentration of exposure. Second, we show cell morphology is a strong indicator of response to differential PM exposure. Third, we used single cell clones to show that specific starting morphologies can encode susceptibility to air pollution exposure. Collectively, our findings show that cell morphology has the potential to be used as a biomarker for environmental risk assessment (Fig. 7).

**Figure 7.**
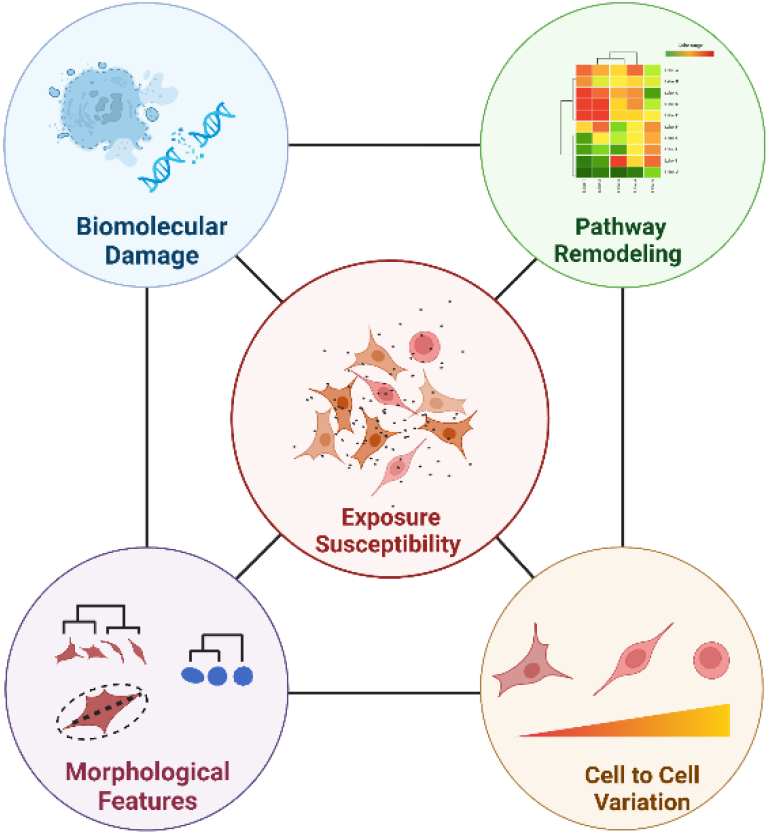
Morphology encodes susceptibility and is dependent upon the interplay between molecular changes in cells.

With the development of single-cell technologies, in both transcriptomics and morphological contexts, and advances in RNA fluorescence *in situ* hybridization (FISH) techniques, further studies could be performed to more directly link the expression of different transcripts with morphological features of individual cells. Additionally, the use of primary cells in a more realistic extracellular matrix environment and the testing of additional PM mixtures could further improve the biological context of future work. Lastly, the use of live-cell imaging to monitor cellular changes over time could lead to a better understanding of stability of morphological patterns which would help better understand susceptibility.

Taken together, our data begins to elucidate how different PM mixtures drive unique changes in morphological and transcriptional signatures, and individual cells within a population have differing levels of susceptibility, encoded for in their morphologies. This knowledge could provide a better understanding of how components of particulate matter such as cadmium and other metals drive PM toxicity. Furthermore, our findings could facilitate the development of a morphology-based method for characterizing an individual’s risk to air pollution exposure.

## Materials and Methods Cell

### Culture

BEAS-2B cells (ATCC CRL-9609) were cultured from cryopreserved stocks in collagen-coated T-75 culture flasks according to ATCC guidelines. Briefly, cells were seeded at 3,000 cells/cm^2^ and cultured in 23mL of BEGM Bronchial Epithelial Cell Growth Medium (Lonza, CC-3170), omitting the addition of the gentamicin-amphotericin aliquot to the medium, as recommended by ATCC. Cells were grown at 37°C in a humidified incubator with a 5% CO_2_ atmosphere, and complete media exchanges were performed every 48 hours. After approximately 4 days, the cultures reached ∼70% confluency, and cells were sub-cultured into 6-well or 96-well plates coated with Type 1 collagen (Advanced BioMatrix, Cat#5005), and allowed to attach to the growth surface for 24 hours prior to exposure to particulate matter.

### Particulate Matter Exposure

The BEAS-2B cells were exposed to three particulate matter mixtures collected from different sources that were purchased from NIST, Urban Particulate Matter (SRM 1648a), Fine Atmospheric Particulate Matter (SRM 2786), and Diesel Exhaust Particulate Matter (SRM 2975). Just prior to the start of the exposures, the three PM mixtures were weighed using an analytical balance and suspended in sterile DI H2O in 10mg/mL stock solutions. The suspensions were sterilized by UV irradiation for 30 minutes as done previously(40). Serial dilutions were performed with BEGM medium to reach the tested concentrations between 1000µg/mL and 31µg/mL. For exposures that contained supplements of cadmium, cadmium chloride was dissolved in DI H_2_O and filter sterilized prior to being added to the PM mixtures at 1000x dilutions. To begin the exposures, media from the well plates was removed and replaced with equal volumes PM-containing media for exposed cells or fresh media for unexposed control cells. The cells were incubated at 37°C in a humidified incubator with a 5% CO_2_ atmosphere for a 24-hour exposure period prior to downstream analysis.

### AlamarBlue**™** assay

BEAS-2B cells were seeded in 96 well plates at a density of 10,000 cells/well. 24 hours later, the cells were exposed to PM mixtures at concentrations ranging from 31-1000 µg/mL as described above with n=7 replicates per condition. Following a 24h exposure period, the media containing the PM was removed and 100µL of fresh BEGM containing 10% alamarBlue™ (Invitrogen, DAL1025) by volume was added to each well. Cells were then incubated at 37°C for 2 hours in the dark. Following this incubation, the fluorescence of each well was measured (Ex. 560/Em. 590) using a BioTek Cytation3 microplate reader. The fluorescence readouts correspond to cell metabolic activity and were normalized to the readings from unexposed control cells after performing background correction by subtracting the fluorescence of wells containing only the alamarBlue™-BEGM mixture.

### Annexin V – Propidium Iodide Flow Cytometry

In this assay, levels of the FITC-labeled Annexin V protein indicate apoptosis as the A.V protein binds with high affinity to the phosphatidylserine that is translocated from the inner side of the cell membrane to the outer side. Likewise, levels of propidium iodide (P.I.), which fluoresces upon binding DNA in cells that have ruptured or become permeable, indicate cell death or cells that are in the latest stages of apoptosis(55, 56). The preparation of cells for flow cytometry was conducted according to established protocols(55). Briefly, following the completion of PM exposures using n=3 replicates, culture media was collected and put on ice to recover detached cells. Adherent cells were trypsinized and combined with the collected culture media. The combined cells were washed twice with cold PBS before proceeding with Annexin V-FITC and propidium iodide staining of 250,000 cells per sample using an eBioscience™ Annexin V Apoptosis Detection Kit (ThermoFisher, 88-8005-72). Prepared samples were analyzed on a Sony Biotechnology MA900 Cell Sorter available through the Center for Biomedical Research Support at UT Austin. At least 10,000 cells per replicate were analyzed for Annexin V binding and propidium iodide incorporation.

### Cell Staining and Imaging

Following exposure, cells adhered to cover glass coated with Type 1 collagen (Advanced BioMatrix, Cat#5005) were washed with prewarmed PBS for 5 minutes then fixed by incubation for 15 minutes at 37°C in a freshly prepared, methanol-free 4% formaldehyde solution in PBS. Cells were rinsed 3x with PBS before being permeabilized by incubation in a 0.1% Triton-X PBS solution for 4 min. Cells were again rinsed 3x with PBS and then blocked with 1% BSA in PBS for 20 minutes at room temperature. Cells were incubated with a 1:400 dilution of a recombinant anti-γH2AX (phosphoS139) antibody (Abcam, ab81299) overnight at 4°C to visualize the DNA damage biomarker. The next day, cells were washed 3x with PBS for 5 min and then incubated with a 1:250 dilution of a fluorescently-tagged secondary antibody (Goat Anti-Rabbit IgG H&L (Alexa Fluor® 488) (Abcam, ab150077)) for 1h at RT. Cells were then rinsed 3x with PBS and stained with Alexa Fluor™ 594 Phalloidin (Invitrogen, A12381) and Invitrogen™ NucBlue™ Fixed Cell ReadyProbes™ Reagent (DAPI) (Invitrogen, R37606) according to the manufacturers’ protocols to allow visualization of the F-actin structure and nuclei, respectively. Microscopy slides were then assembled using ProLong™ Gold Antifade Mountant (Invitrogen, P36930) and were sealed with clear nail polish. Slides were stored at 4°C until imaging.

Fluorescent images were acquired with a Leica Stellaris 5 Confocal Microscope at 20X resolution using 3 laser lines (405 Diode: DAPI Nuclear Stain, 488 Diode: Alexa Fluor® 488 secondary antibody targeting γH2AX, 647 Diode: Phalloidin/Actin Stain). Individual Nuclei/Cell Boundaries were segmented with Cell Profiler^TM^ (57) in combination with in-house curation pipelines to ensure well-segmented single cells. Briefly, an immunofluorescence-focused segmentation algorithm used the DAPI stain to delineate the nucleus shape and the Phalloidin stain to delineate the general cell shape. Approximately 13,000 single cells spanning all exposure conditions were procured for this work with an additional 40,000 cells analyzed for single cell clones.

### Data Processing and Morphological Analysis

33 key morphological parameters were extracted from each individual cell using a Cell Profiler^TM^ morphological analysis pipeline (**Table S1**). In order to compare morphological parameters of different scales to understand population variance, all morphological parameters were independently log normalized. This “normalized” morphological parameter dataset was subsequently used to construct a 2D-Uniform Manifold and Projection (UMAP) space(58). UMAP is a nonlinear dimensionality reduction algorithm that seeks to capture the structure of high dimensional data in a lower-dimensionality space (for this work, the 33-vector space was simplified down to two). Each point in the UMAP space represents an individual cell whose morphological parameters have been transformed and projected onto the 2D-UMAP space. k-means clustering, an unsupervised clustering method, was used to identify distinct morphological groups within the cell dataset from the log-normalized dataset. An optimal number of clusters, ten, was calculated by a plateau in the inertia and silhouette values of the k-means algorithm (**Fig. S13**). To quantify morphological heterogeneity, the Shannon entropy for each PM exposure condition was calculated using the k-means clusters as follows.

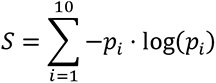

Where *S* is the Shannon entropy (greater magnitude signifies a more heterogeneous population) and *p_i_* is the fraction of the population that is in morphological cluster i(52). For single cell cloning analysis, larger morphological cluster groups were created to identify overarching morphological themes of the k-means clusters. Briefly, ward-based clustering was performed on the average morphological signature across each k-means cluster, and the analysis identified 3 morphological groups that encompassed the k-means clusters.

γH2AX content per cell was analyzed through the mean nuclear intensity of the fluorescent 488 channel. Specifically, the summation of the pixel values (normalized to range from 0-1) of the 488 channel was divided by the pixel area of the encompassing nuclei. The resulting mean γH2AX expression was then layered across the UMAP manifold and analyzed per cluster. Approximately 13,000 individual cells encompassing all PM exposures were analyzed for the morphological analysis.

### Single Cell Cloning and Live-Cell Imaging

Single cells of the BEAS-2B cell line were isolated using a Sony Biotechnology MA900 Cell Sorter available through the Center for Biomedical Research Support at UT Austin. Individual cells were sorted into a 96-well plate and allowed to proliferate. Media exchanges of BEGM were performed every 48 hours. Cell populations were expanded to collagen coated 24-well plates, 6-well plates, and finally T-75 flasks before freezing cells to create multiple clonal populations. Clonal populations were then similarly used in experiments as the parental BEAS-2B population as described above. Approximately 40,000 single cells spanning all clones and urban exposure conditions were analyzed.

### 3’-Tag RNA Sequencing

BEAS-2B cells were cultured and exposed to PM as described above. Following the completion of 24h exposures to the three PM types at two concentrations (125 and 500 µg/mL), RNA extraction was immediately performed on n≥4 replicates by lysing cells with TRIzol™ Reagent (Invitrogen, 15596026). The RNA underwent DNase I treatment and was purified using a Direct-zol RNA Miniprep Kit (Zymo Research, R2052) according to manufacturer protocol. The purity of the RNA was confirmed using a Nanodrop 2000 Spectrophotometer (Thermo Scientific), and RNA concentration was determined using a Qubit™ 4 Fluorometer (ThermoFisher) RNA Broad Range Assay Kit (ThermoFisher, Q10210). Prior to library preparation, RNA quality was determined using an Agilent Bioanalyzer and all samples used for sequencing had a RIN score >8.80. The RNA was submitted to the University of Texas Genomic Sequencing and Analysis Facility for 3’ RNA based library preparation and sequencing based on previously published protocols(41, 42). Libraries were quantified using the Quant-it PicoGreen dsDNA assay (ThermoFisher) and pooled equally for subsequent size selection at 350-550bp on a 2% gel using the Blue Pippin (Sage Science). The final pools were checked for size and quality with the Bioanalyzer High Sensitivity DNA Kit (Agilent) and their concentrations were measured using the KAPA SYBR Fast qPCR kit (Roche). The pooled libraries were sequenced on a NovaSeq6000 (Illumina) and a sequencing depth of 4.5 million reads per sample was achieved with single-end, 100-bp read length. Raw sequencing data is available at the National Center for Biotechnological Information (NCBI) Short Read Archive (SRA) under BioProject Accession no. PRJNA954385.

### Differential Gene Expression Analysis

Following sequencing, the raw reads were preprocessed to remove adapter contamination and trim the unique molecular identifier (UMI) barcodes, remove duplicates, and remove poor quality reads. The Human Reference Genome was assembled and indexed using Homo_sapiens.GRCh38.dna.primary_assembly.fa and Homo_sapiens.GRCh38.104.gtf from Ensembl using the genomeGenerate run mode in STAR(59). The filtered reads were then aligned to the generated genome using STAR. HTSeq(60) was used to count the aligned reads in each .bam file generated by STAR. The DESeq2 package(61) was then used to quantify differential gene expression in R(62). Differential expression was determined for each PM exposure condition relative to the counts from unexposed control cells. Significantly differentially expressed genes were defined as having a log_2_(Fold Change)≥1 and p_adj_<0.05. GO Term analysis was performed using the Enrichr web tool(45) to determine GO Biological Process terms that were significantly enriched in the sets of significantly differentially expressed genes. Significant GO Terms were defined as having p_adj_<0.01. Chord plots were constructed using the GOplot package in R.

## Acknowledgments

We acknowledge the funding support of this study from the National Institutes of Health R21ES032124 (L.M.C.) and U01AG060903 (J.M.P.), the National Science Foundation EF-2022124 (L.M.C.), and The Johns Hopkins University Older Americans Independence Center of the National Institute on Aging (NIA) under award number P30AG021334 (JMP). This work is supported by the National Science Foundation Graduate Research Fellowship Program under Grant No. DGE 2137420 (S.M.E.). Any opinions, findings, and conclusions or recommendations expressed in this material are those of the authors and do not necessarily reflect the views of the National Science Foundation. The authors acknowledge the Texas Advanced Computing Center (TACC) at UT Austin for providing high performance computing resources. We also thank the Genomic Sequencing and Analysis Core Facility at UT Austin, RRID#: SCR_021713, for assistance with RNA-sequencing.

