## Supplemental Figures for "Particulate matter composition drives differential molecular and morphological responses in lung epithelial cells"

**This PDF file includes:**

Figures S1 to S13
Table S1

**Other supporting materials for this manuscript include the following:**

Datasets S1 to S13

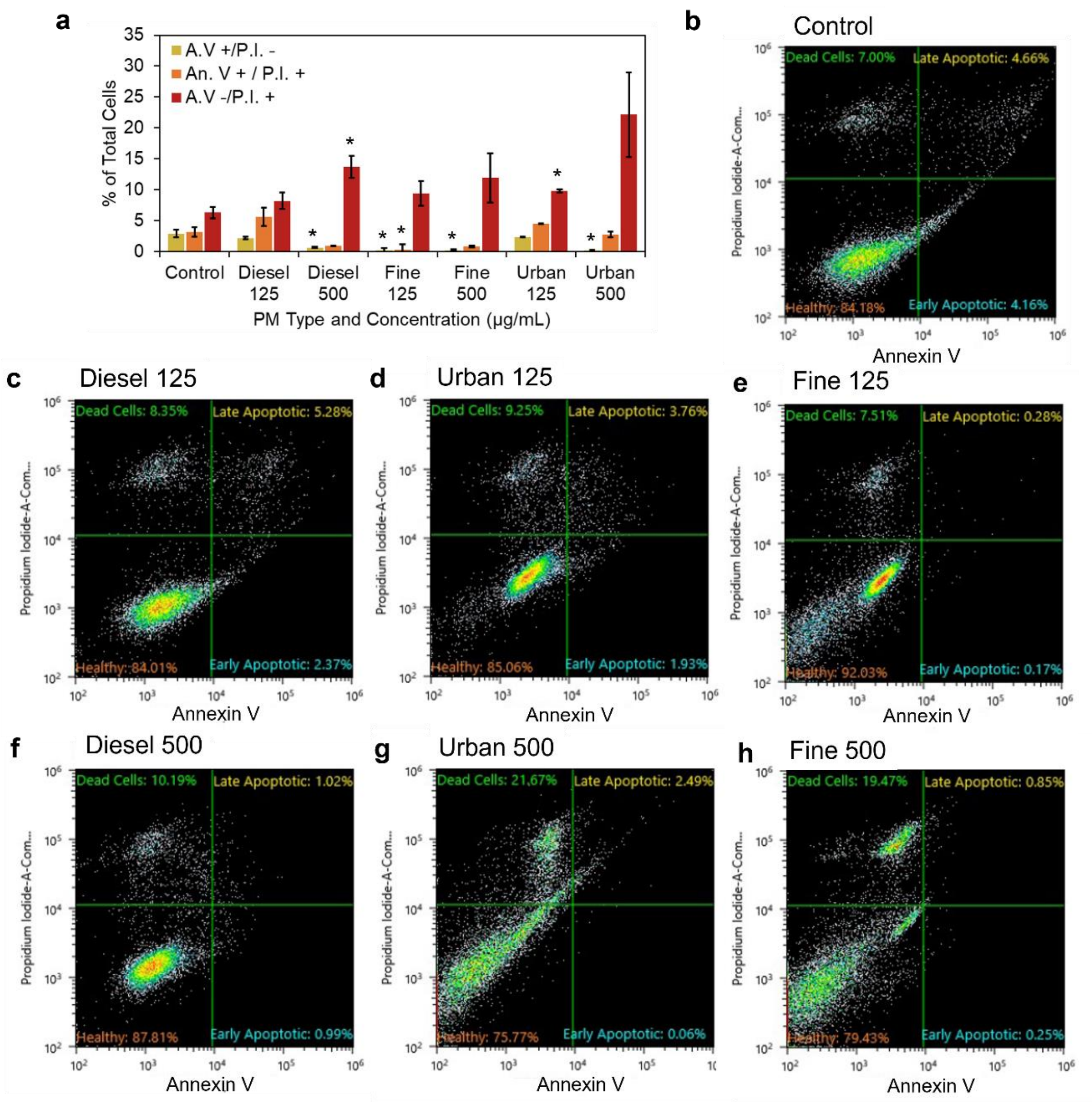


**Fig. S1.** (**a**) Flow cytometry analysis of Annexin V-Propidium Iodide apoptosis assay following PM exposure. A.V+/P.I.- and A.V+/P.I.+ represent early- and late-stage apoptotic cells respectively, A.V-/P.I.+ represents dead cells (n=3, ≥10,000 cells per measurement, error bars represent the standard error of the mean). (**b-h**) Representative flow cytometry scatter plots for analysis of cells following 24h exposure to different PM types and concentrations (500 or 125 µg/mL). displaying the distribution of cells according to Annexin V-FITC intensity (x axis) and propidium iodide intensity (y axis).

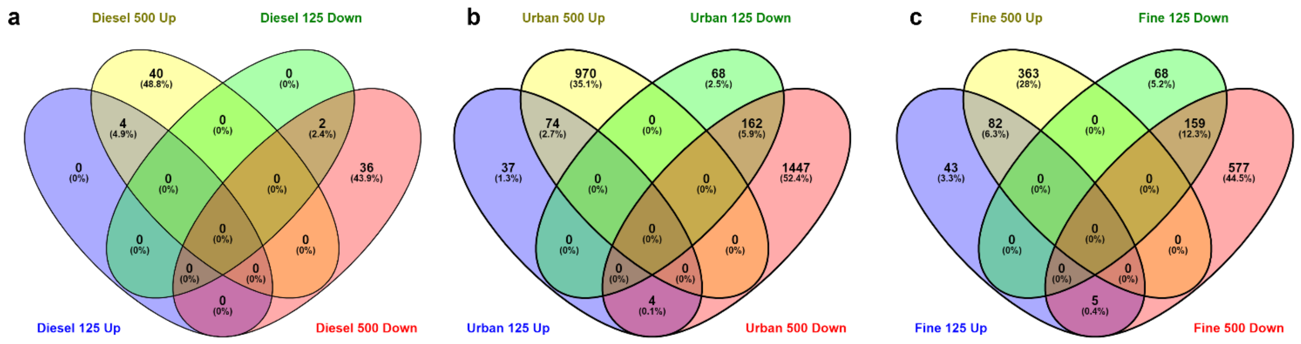


**Fig. S2.** Venn diagrams of genes differentially upregulated (Up) and downregulated (Down) at the 125µg/mL and 500µg/mL concentrations for (**a**) Diesel Exhaust, (**b**) Urban, and (**c**) Fine PM exposures. The majority of up and down regulated genes after the 125µg/mL exposures continue to be up and down regulated after the 500µg/mL exposures respectively.

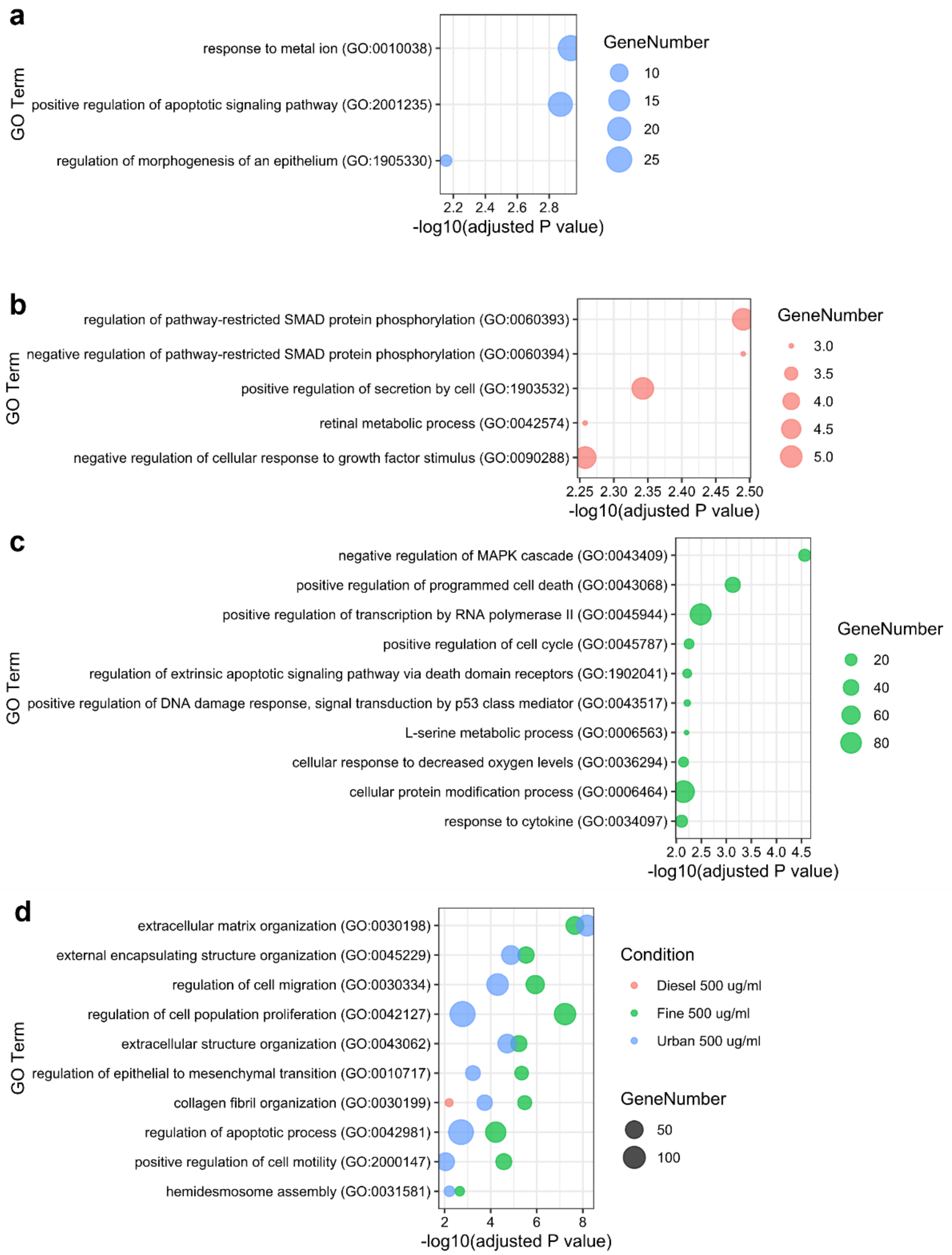


**Fig. S3.** Enriched GO Biological Process terms resulting from high level (500 µg/mL) exposures. Significantly enriched terms ($p_{adj}<0.01$) that are unique to each individual condition are shown in (**a-c**): (**a**) Urban, (**b**) Diesel, (**c**) Fine. Terms that are shared among two or more conditions and represent a common response are shown in (**d**). Note, no additional terms were significant and unique to the Urban or Diesel conditions. Complete GO analysis can be found in Supplemental Datasets S8-13.

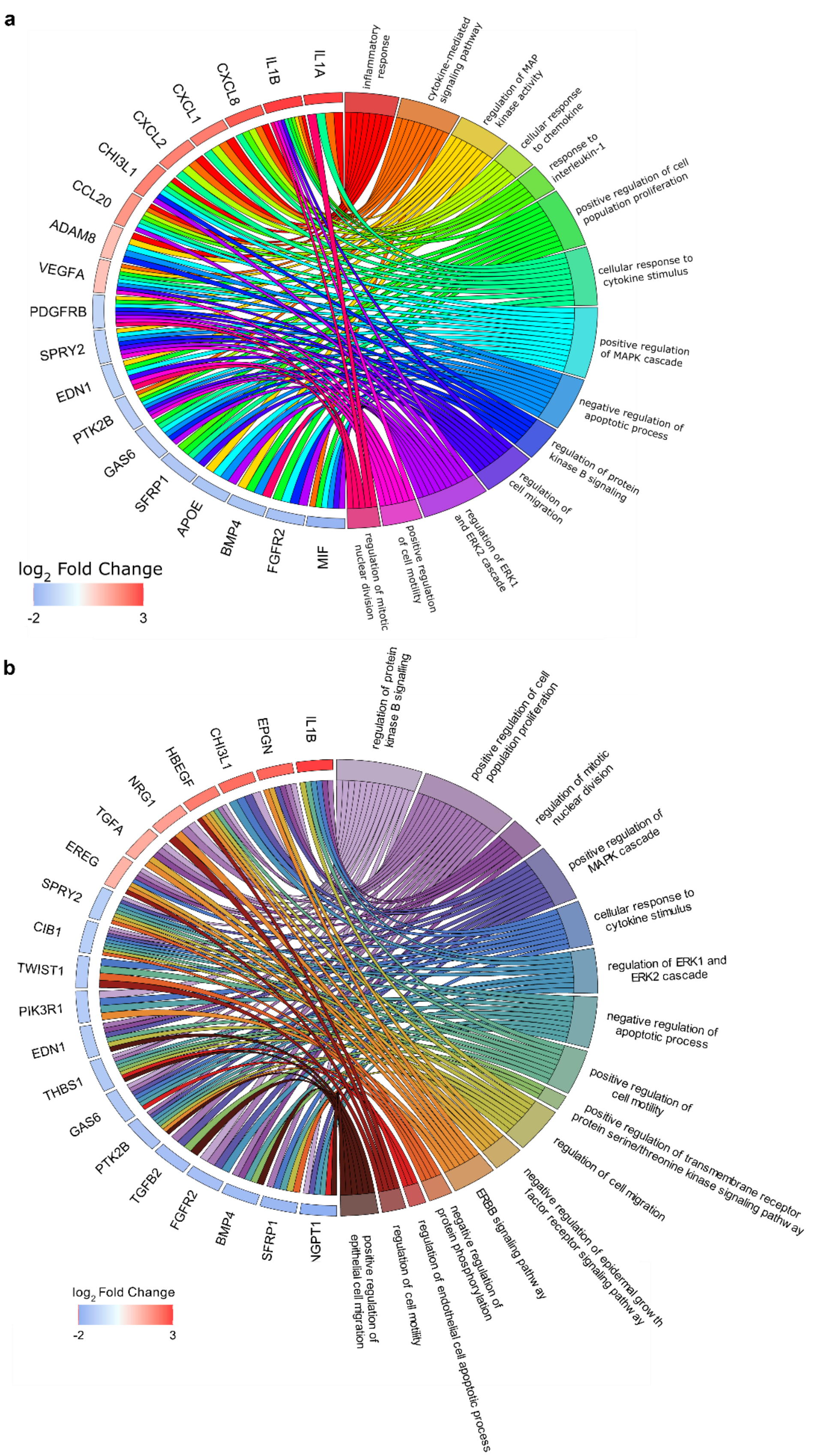
a
IL1A
IL1B
CXCL8
CXCL1
CXCL2
CHI3L1
CCL20
ADAM8
VEGFA
PDGFRB
SPRY2
EDN1
PTK2B
GAS6
SFRP1
APOE
BMP4
FGFR2
MIF
inflammatory response
cytokine-mediated signaling pathway
regulation of MAP kinase activity
cellular response to chemokine
response to interleukin-1
positive regulation of cell population proliferation
cellular response to cytokine stimulus
positive regulation of MAPK cascade
negative regulation of apoptotic process
regulation of protein kinase B signaling
regulation of cell migration
regulation of ERK1 and ERK2 cascade
positive regulation of cell motility
regulation of mitotic nuclear division
log2 Fold Change
-2
3
b
IL1B
EPGN
CHI3L1
HBEGF
NRG1
TGFA
EREG
SPRY2
CIB1
TWIST1
PIK3R1
EDN1
THBS1
GAS6
PTK2B
TGFB2
FGFR2
BMP4
SFRP1
NGPT1
regulation of protein kinase B signaling
positive regulation of cell population proliferation
regulation of mitotic nuclear division
positive regulation of MAPK cascade
cellular response to cytokine stimulus
regulation of ERK1 and ERK2 cascade
negative regulation of apoptotic process
positive regulation of cell motility
positive regulation of transmembrane receptor protein serine/threonine kinase signaling pathway
regulation of cell migration
negative regulation of epidermal growth factor receptor signaling pathway
ERBB signaling pathway
negative regulation of protein phosphorylation
regulation of endothelial cell apoptotic process
regulation of cell motility
positive regulation of epithelial cell migration
log2 Fold Change
-2
3

**Fig. S4.** Chord plots displaying the relationships between enriched GO Terms and the associated genes for the 125µg/mL Fine (**a**) and Urban (**b**) exposure conditions. Genes are listed in descending order of differential expression, and the magnitude of gene differential expression is displayed as a color scale of the $Log_2$ Fold Change value next to each gene name. A connection between a gene and pathways indicates the gene is involved in that pathway. Genes involved in fewer than 4 pathways are omitted, and pathways with fewer than 3 differentially expressed genes are omitted.

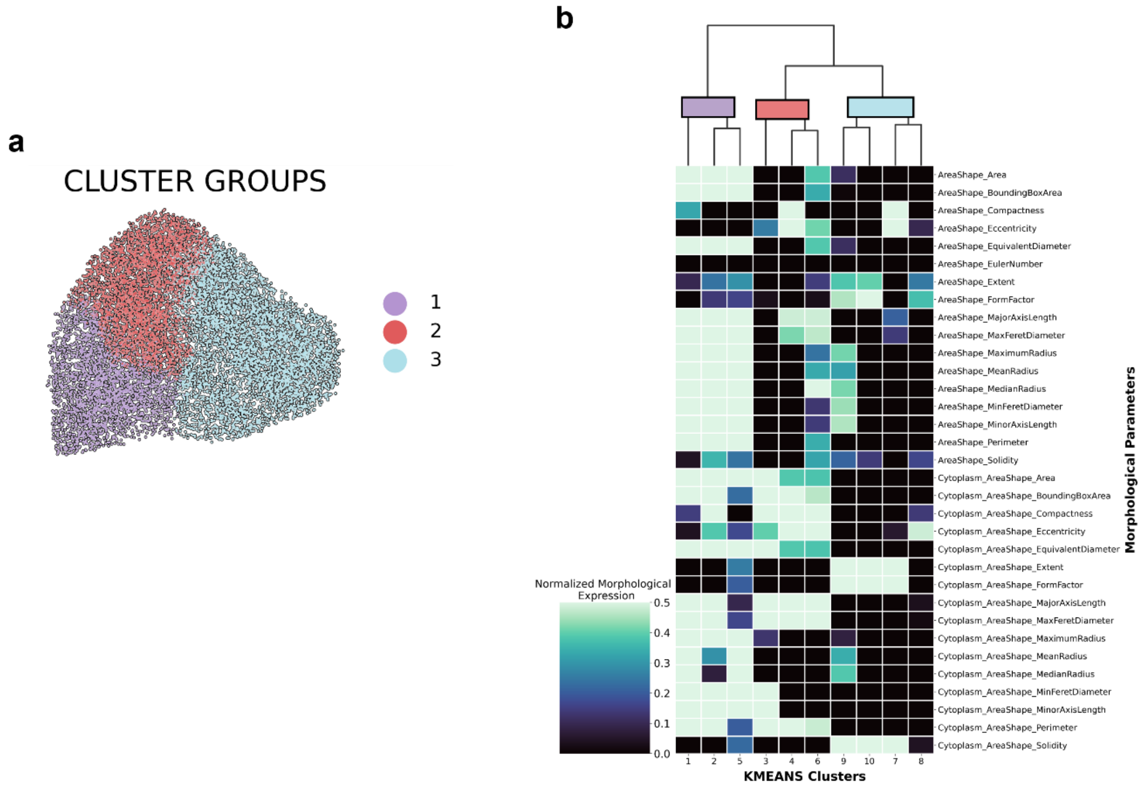


**Fig. S5.** (**a**) The 10 k-means clusters that are used to define cell morphology can be further grouped into 3 cluster groups (CG1-3) using Ward based clustering. (**b**) Dendrogram displaying how the clusters were grouped using the 33 measured morphological parameters.

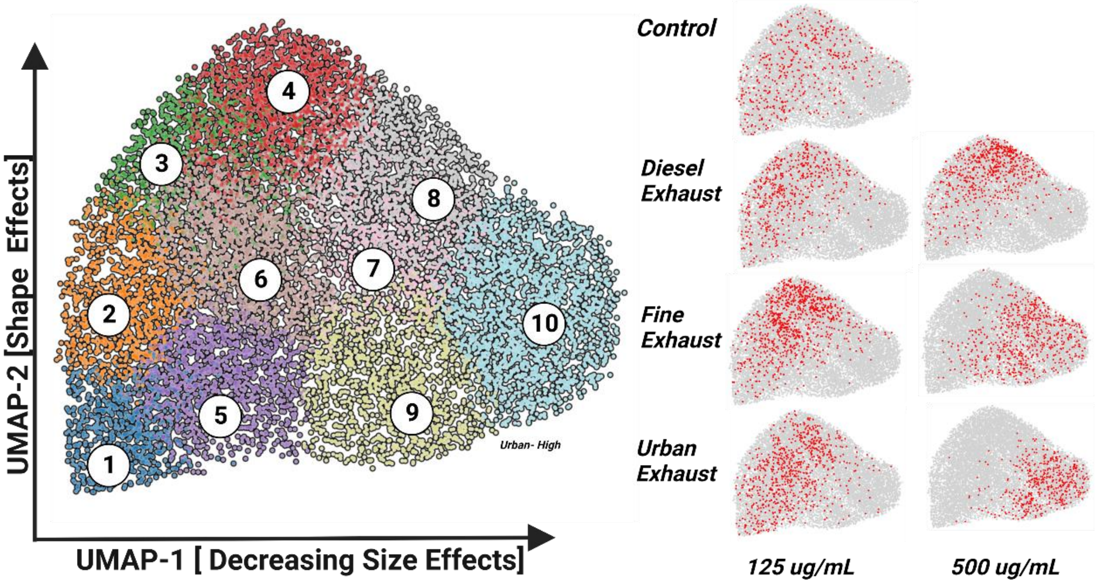


**Fig. S6.** UMAP visualization of the 33 measured morphological parameters for each cell in every condition. UMAP-1 (X-axis) was *negatively* correlated with size and UMAP-2 (Y-axis) was *positively* correlated with cell elongation, or linearity. K-means clustering was applied to cluster cells of similar morphologies. The plots on the right show the distribution of cells from each respective exposure condition in red in the UMAP space.

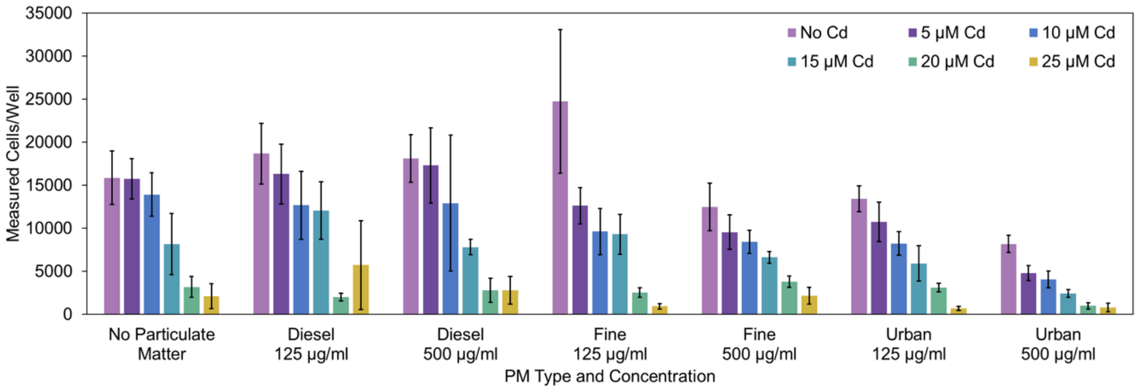


**Fig. S7.** Changes in viable cell counts as a result of PM and cadmium supplementation exposure. Cell counting was performed using the Trypan Blue exclusion method. A hemocytometer was used for counting n=4 replicates, error bars represent one standard deviation.

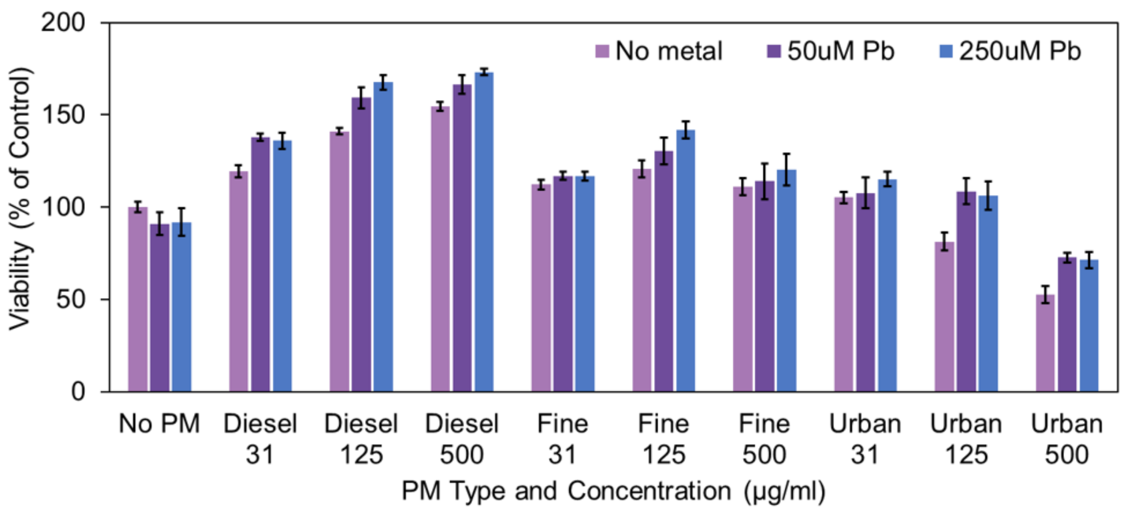


**Fig. S8.** Cell viability following 24h exposures to different PM types and concentrations supplemented with lead acetate (0-250 µM Pb). Values are percentages of viable cells relative to unexposed control cells as measured with the alamarBlue assay (n=6, error bars represent one standard deviation).

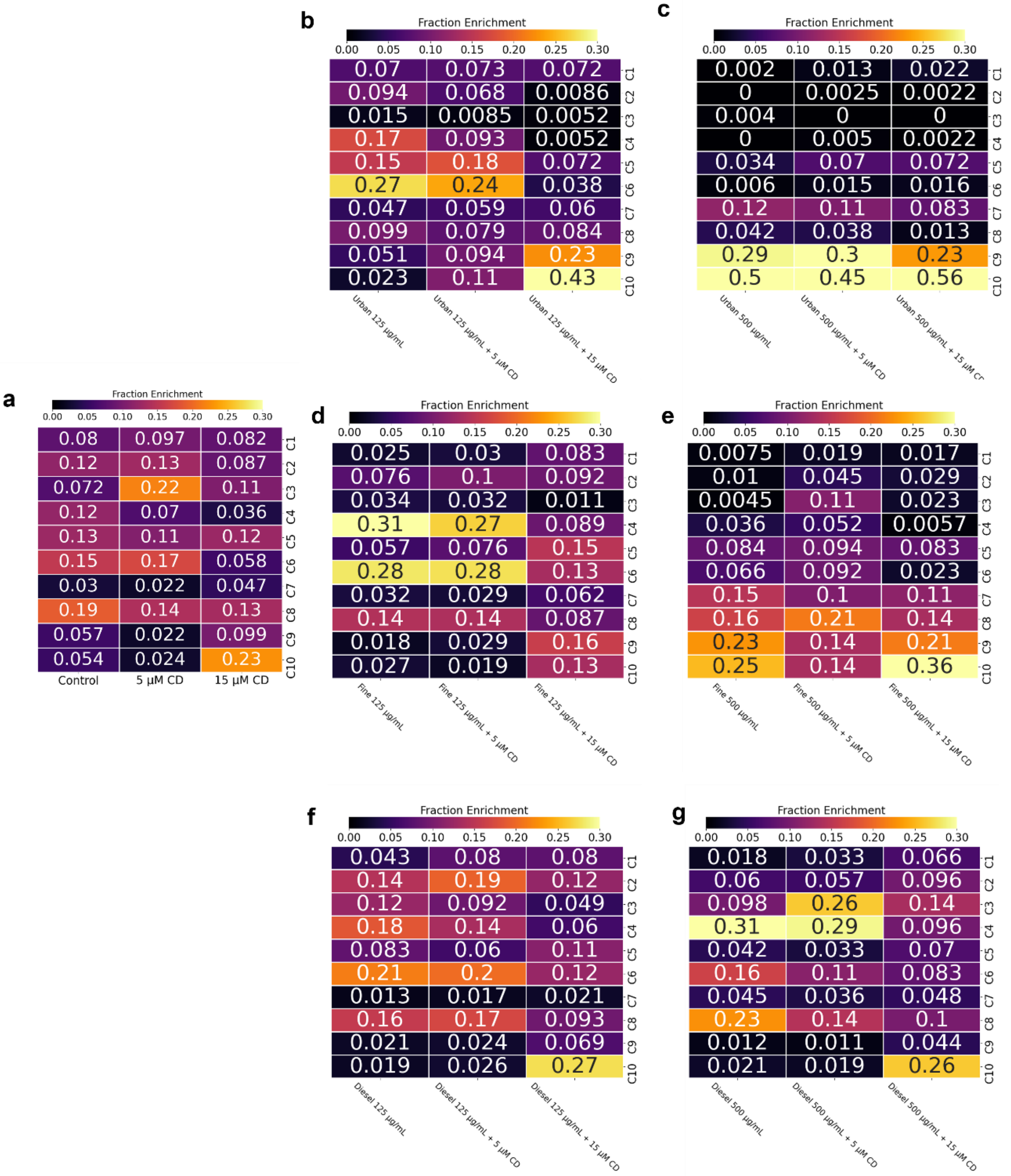


**Fig. S9.** Heatmaps displaying the fractions of cells across the 10 k-means morphological clusters for each PM exposure condition with cadmium supplementation.

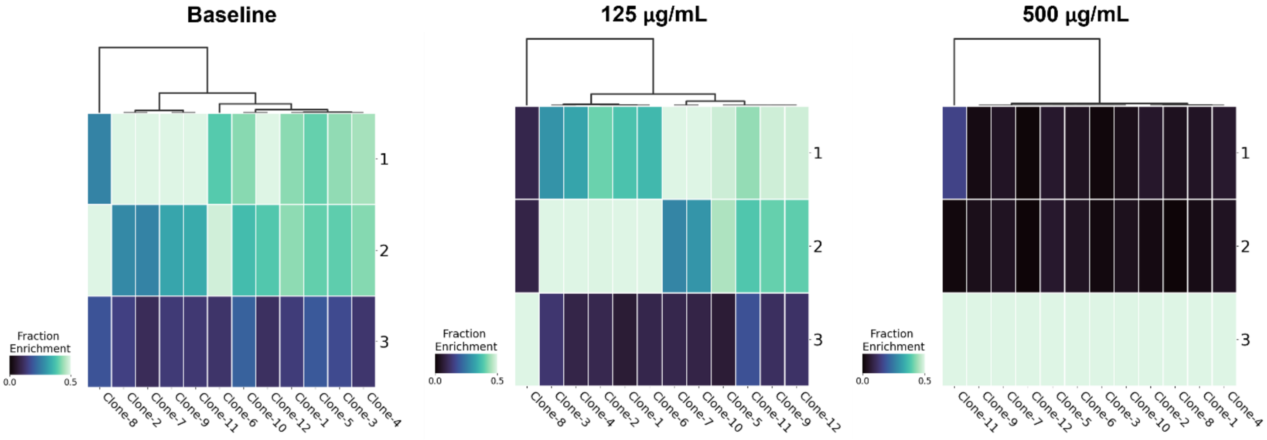


**Fig. S10.** Heatmaps displaying the enrichment in number of cells in the three morphology cluster groups for each clonal population. Each heatmap represents morphological distributions from different conditions, either (**a**) baseline morphology, or morphology following the (**b**) 125µg/mL or (**c**) 500µg/mL Urban PM exposure.

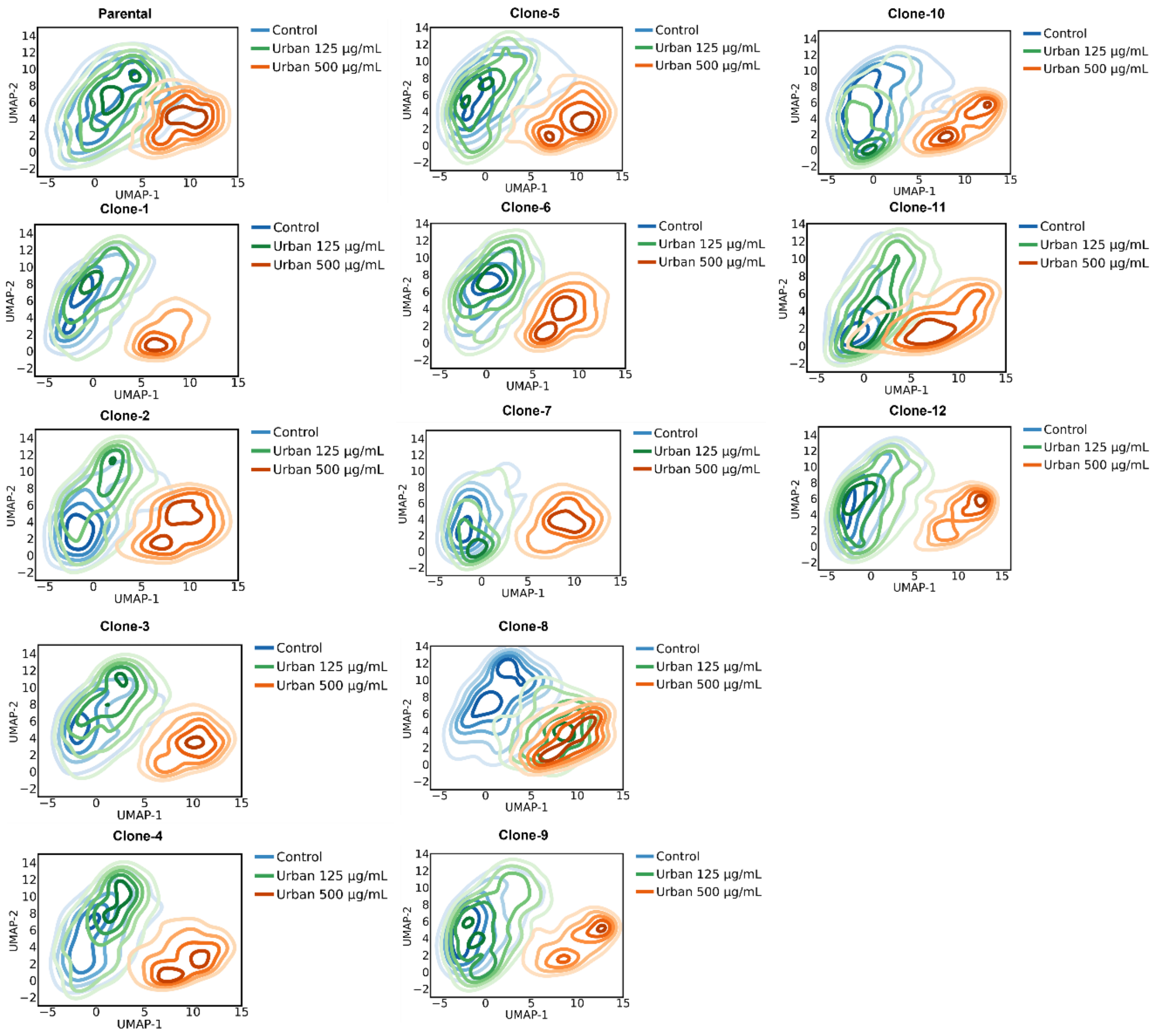


**Fig. S11.** Contour maps showing the distributions of morphologies of cells for each clonal population across the UMAP space.

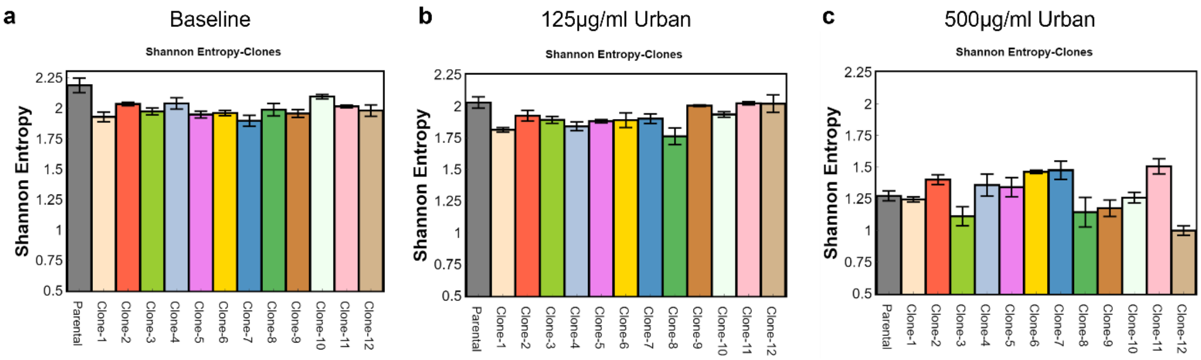


**Fig. S12**. The Shannon Entropies are lowered at baseline for each single cell clone population relative to the parental population. Shannon entropies are shown for (**a**) the control unexposed condition, (**b**) post-exposure to 125µg/ml Urban PM, and (**c**) post-exposure to 500ug/ml Urban PM. Error bars represent standard error of the mean.

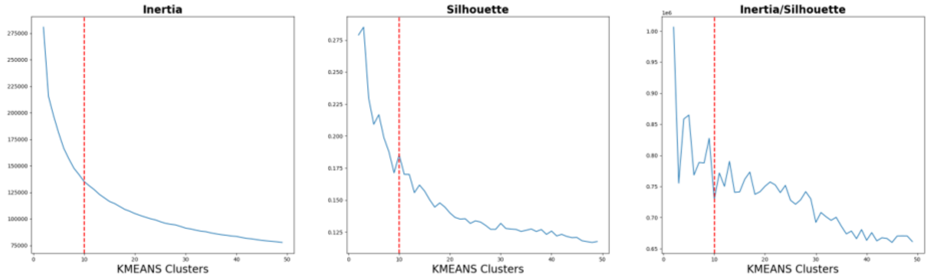


**Fig. S13.** Inertia and silhouette values calculated for various numbers of k-means clusters. The optimal number of clusters, 10, was selected for its local minimum in its inertia/silhouette value and for expressing a reasonable number of clusters.

**Table S1.** List of the morphological parameters that were quantified for each cell in the morphological analysis.

| Nuclear | Cellular |
|---|---|
| Nuclear Area | Cellular Area |
| Nuclear Bounding Box Area | Cellular Bounding Box Area |
| Nuclear Compactness | Cellular Compactness |
| Nuclear Eccentricity | Cellular Eccentricity |
| Nuclear Equivalent Diameter | Cellular Equivalent Diameter |
| Nuclear Euler Number | Cellular Extent |
| Nuclear Extent | Cellular Form Factor |
| Nuclear Form Factor | Cellular Major Axis Length |
| Nuclear Major Axis Length | Cellular Maximum Feret Diameter |
| Nuclear Maximum Feret Diameter | Cellular Maximum Radius |
| Nuclear Maximum Radius | Cellular Mean Radius |
| Nuclear Mean Radius | Cellular Median Radius |
| Nuclear Median Radius | Cellular Minimum Feret Diameter |
| Nuclear Minimum Feret Diameter | Cellular Minor Axis Length |
| Nuclear Minor Axis Length | Cellular Periometer |
| Nuclear Periometer | Cellular Solidity |
| Nuclear Solidity | |
